# Immune surveillance in clinical regression of pre-invasive squamous cell lung cancer

**DOI:** 10.1101/833004

**Authors:** Adam Pennycuick, Vitor H. Teixeira, Khalid AbdulJabbar, Shan E Ahmed Raza, Tom Lund, Ayse Akarca, Rachel Rosenthal, Christodoulos P. Pipinikas, Henry Lee-Six, Deepak P. Chandrasekharan, Robert E. Hynds, Kate H. C. Gowers, Jake Y. Henry, Celine Denais, Mary Falzon, Sophia Antoniou, Pascal F. Durrenberger, Andrew Furness, Bernadette Carroll, Ricky M. Thakrar, Philip J. George, Teresa Marafioti, Charles Swanton, Christina Thirlwell, Peter J. Campbell, Yinyin Yuan, Sergio A. Quezada, Nicholas McGranahan, Sam M. Janes

## Abstract

Before squamous cell lung cancer develops, pre-cancerous lesions can be found in the airways. From longitudinal monitoring, we know that only half of such lesions become cancer, whereas a third spontaneously regress. While recent studies have described the presence of an active immune response in high-grade lesions, the mechanisms underpinning clinical regression of pre-cancerous lesions remain unknown. Here, we show that host immune surveillance is strongly implicated in lesion regression. Using bronchoscopic biopsies from human subjects, we find that regressive carcinoma in-situ lesions harbour more infiltrating immune cells than those that progress to cancer. Moreover, molecular profiling of these lesions identifies potential immune escape mechanisms specifically in those that progress to cancer: antigen presentation is impaired by genomic and epigenetic changes, TGF-beta signalling is overactive, and the immunomodulator TNFSF9 is downregulated. Changes appear intrinsic to the CIS lesions as the adjacent stroma of progressive and regressive lesions are transcriptomically similar. This study identifies mechanisms by which pre-cancerous lesions evade immune detection during the earliest stages of carcinogenesis and forms a basis for new therapeutic strategies that treat or prevent early stage lung cancer.

---

Before the development of lung squamous cell carcinoma (LUSC), pre-invasive lesions can be observed in the airways. These evolve stepwise, progressing through mild and moderate dysplasia (low-grade lesions) to severe dysplasia and carcinoma in-situ (CIS; high-grade lesions), before the development of invasive cancer(1). Markers of immune sensing and escape have been associated with increasing grade(2). However, longitudinal bronchoscopic surveillance of such lesions has shown that progression of pre-invasive lesions to cancer is not inevitable; only half of high-grade CIS lesions will progress to cancer within two years, whereas a third will spontaneously regress(3). Here, we integrate genomic, transcriptomic, epigenetic and imaging data across carefully phenotyped airway CIS lesions and adjacent stroma (Table S1; Extended Data Figure 1) to assess the role of immune surveillance in lesion regression. We identify key immune escape mechanisms enriched in pre-invasive lesions which later progressed to cancer. Understanding these mechanisms may offer new therapeutic strategies to induce regression and prevent the development of invasive disease.

To assess our hypothesis that lesion regression is driven by immune surveillance, we first performed immunohistochemistry (IHC) on 28 progressive and 16 regressive CIS lesions (Figure 1a-b). Regressive lesions showed higher concentrations of intra-lesional cytotoxic CD8+ (p=0.037; Figure 1c) but not CD4+ (p=0.25) or regulatory FOXP3+ (p=0.41) T cells. We then quantified immune cells in stromal regions adjacent to CIS lesions, but found no significant differences between progressive and regressive lesions for CD8+ (p=0.49), CD4+ (p=0.43) or FOXP3+ (p=0.64) cells. We then used a machine-learning approach to quantify lymphocytes from hematoxylin and eosin (H&E) stained slides in a much larger dataset of 113 samples, which similarly contained more infiltrating lymphocytes in regressive lesions (Figure 1c; p=0.023).

**Fig. 1.**
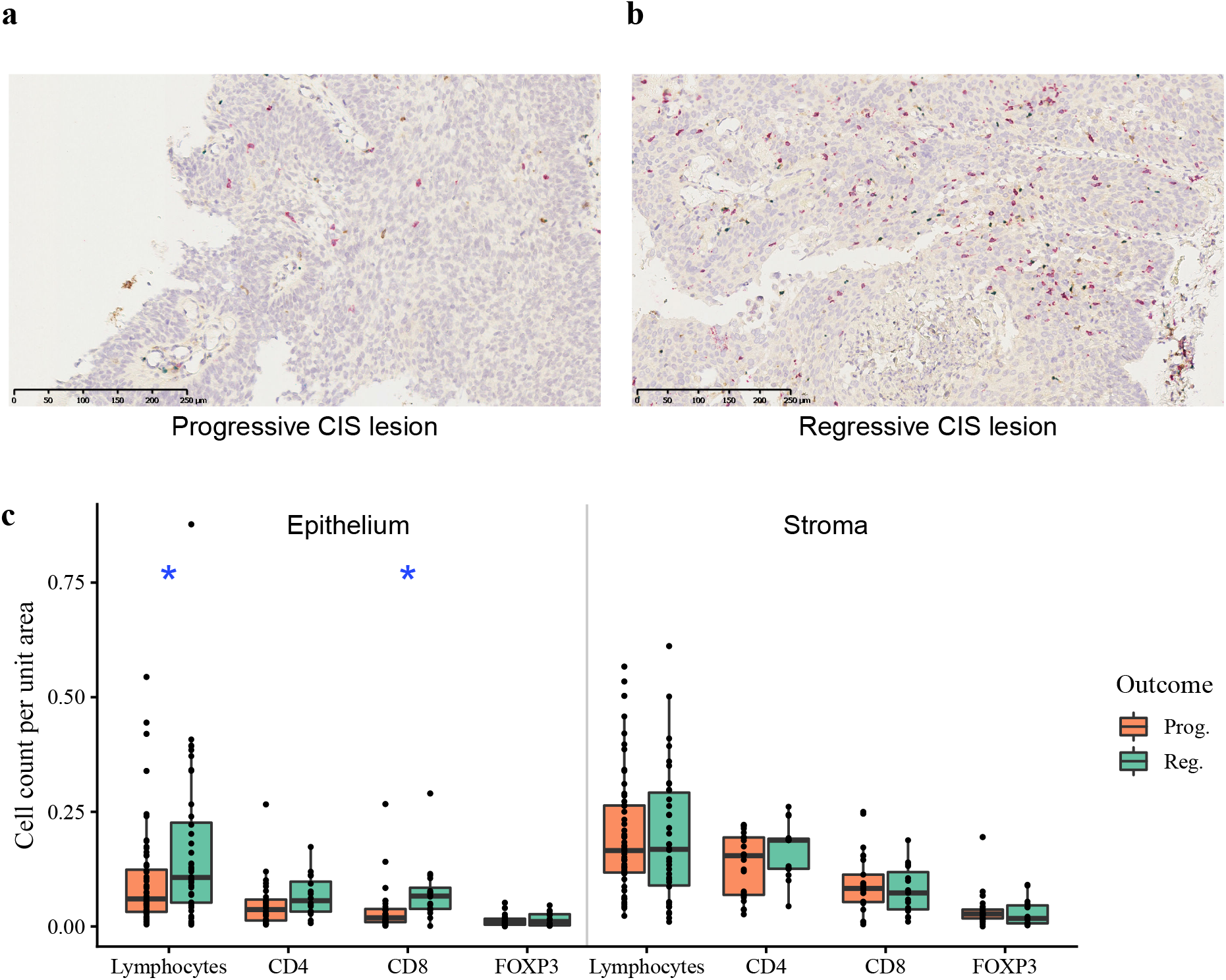
Immune cell infiltration of lung carcinoma-in-situ lesions. (a-b) Immunohistochemistry images of (a) progressive CIS lesion and (b) regressive CIS lesion with CD4+ cells stained in brown, CD8+ cells in red and FOXP3+ in blue. Immune cells are separately quantified within the CIS lesion and in the surrounding stroma. c) Combined quantitative immunohistochemistry data of CD4, CD8 and FOXP3 staining (n=44; 28 progressive, 16 regressive) with total lymphocyte quantification from H&E images (n=116; 69 progressive, 47 regressive) shown. We observe increased lymphocytes (p=0.023) and CD8+ cells (p=0.037) per unit area of epithelium within regressive CIS lesions compared to progressive. Stromal regions adjacent to CIS lesions showed no significant differences in immune cells between progressive and regressive lesions. p-values are calculated using linear mixed effects models to account for samples from the same patient; *<0.05.

For a broader assessment of transcriptomic differences between CIS lesions and their adjacent stroma, we isolated epithelial tissue and paired stroma separately using laser capture microdissection for 10 progressive and 8 regressive CIS lesions. Similarly to IHC data, cell type deconvolution analysis demonstrated higher infiltrating lymphocytes in regressive lesions (Figure 2a; p=0.0012), as did deconvolution of methylation data from 36 progressive and 18 regressive CIS lesions (Figure 2b; p=0.006). Comparing predictions for individual cell types across gene expression and methylation data found an increase in most immune cell types in regressive lesions compared to progressive, with the exception of macrophages – a potentially immunosuppressive cell type – which were more abundant in progressive lesions (p=0.005; Table S2).

**Fig. 2.**
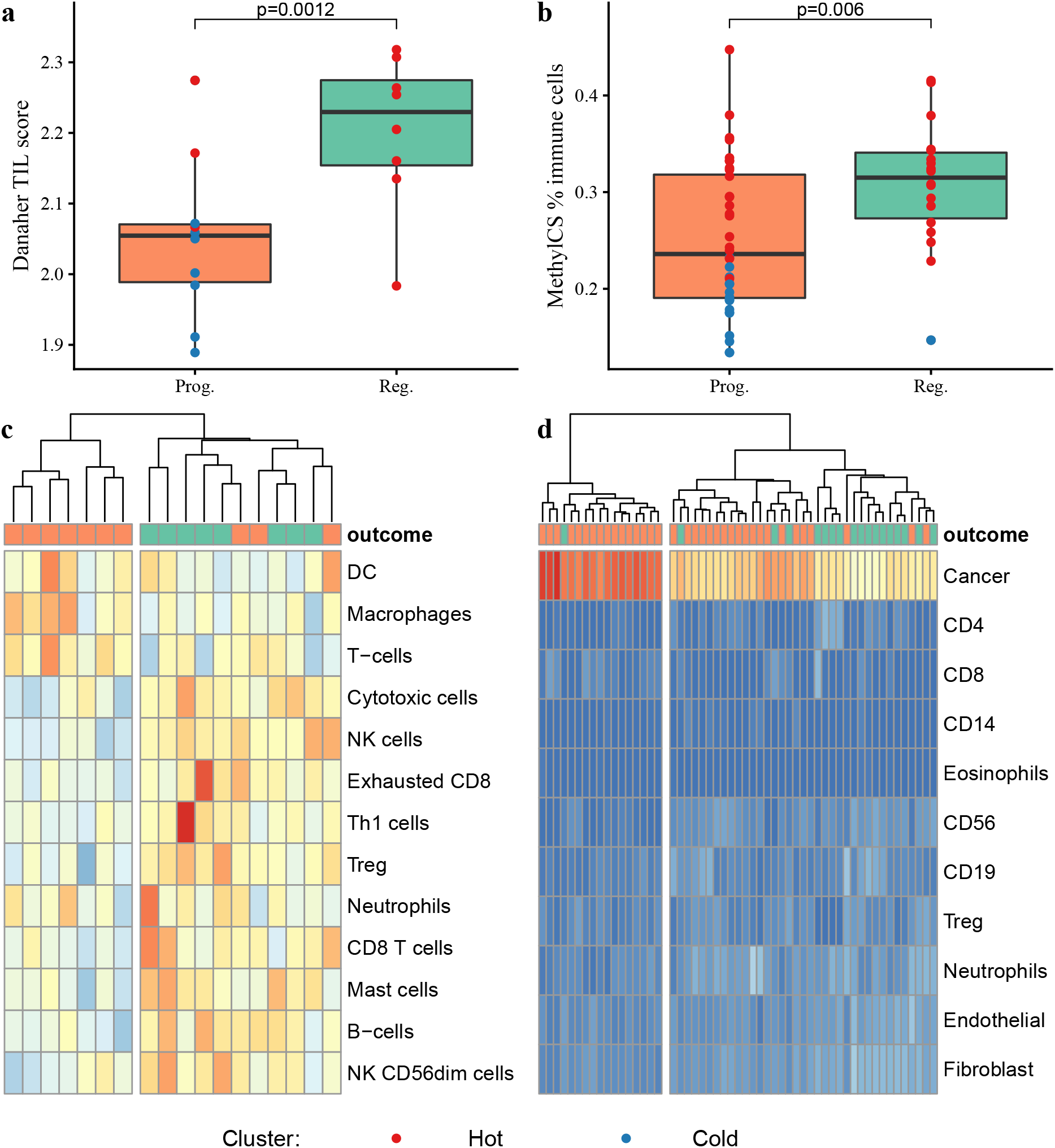
Identification of immune ‘hot’ and ‘cold’ carcinoma in-situ lesions by immune cell clustering. Progressive and regressive lesions have significantly different geneexpression derived TIL scores (a; p=0.0012) and different immune cell percentages as derived from methylCIBERSORT (b; p=0.006). c) Immune cell quantification from gene expression data (n=18) using the method of Danaher et al. shows an ‘immune cold’ cluster (left) in which all lesions progressed to cancer, and an ‘immune hot’ cluster (right) in which the majority regressed. d) Similar clustering on methylation-derived cell subtypes using methylCIBERSORT (n=54) again shows two distinct clusters: an ‘immune cold’ cluster (left) dominated by a cancer cell signature, in which all but one lesion progressed, and an ‘immune hot’ cluster (right), containing both progressive and regressive samples. p-values are calculated using mixed effects models to account for samples from the same patient.

Analysis of proand anti-inflammatory cytokine expression within the epithelial compartment demonstrated an increase in pro-inflammatory (p=1.2×10^−5^) but not anti-inflammatory (p=0.3) response in regressive lesions compared to progressive (Extended Data Figure 2). *IFNG*, *IL2* and *TNF* were all increased in regressive lesions (Extended Data Figure 3). IL10 was also increased in regressive lesions; whilst classically considered an anti-inflammatory cytokine, *IL10* has been shown to stimulate anti-tumor immunity(4). Only *CXCL8* was upregulated in progressive samples compared to regressive (p=1.8×10^−5^); produced by macrophages, the expression of *CXCL8* correlated strongly with macrophage quantification from deconvoluted gene expression data (r^2^=0.62, p=0.007). Taken together, these data are in keeping with a model in which inflammation via IFN-γ, IL-2 and TNF fosters effective immune surveillance, whilst lesion-associated macrophages – similar to tumor-associated macrophages in advanced cancers – have an immunosuppressive effect.

Recent advances have demonstrated heterogeneity of lung cancer immune infiltration, with patients whose tumors have more infiltrated ‘immune hot’ regions having improved survival as compared to those with abundant poorly infiltrated, ‘immune cold’ regions(5, 6). Hierarchical clustering of deconvoluted immune cell quantification at both the transcriptomic and epigenetic levels demonstrated clear clusters of ‘cold’ lesions, almost all of which progressed to cancer (Figure 2c-d). However, we also observed some ‘hot’ progressive lesions, suggesting the presence of other mechanisms in these lesions. We therefore sought to address two questions: firstly, could deficits in antigen presentation and immune recruitment in progressive lesions be identified, which could explain the observed ‘cold’ lesions? Secondly, could disordered immune cell function explain the existence of progressive immune ‘hot’ lesions?

The acquisition of mutations that result in clonal neoantigens drives T cell immunoreactivity in cancer(7). We hypothesised that immune-active regressive lesions may contain more neoantigens than progressive lesions, however, this was not supported by whole-genome sequencing data(8) (n=39). Predicted neoantigens correlated very closely with mutational burden (r^2^=0.94), and progressive lesions have been shown to have significantly higher mutational burden than regressive lesions(8), therefore more neoantigens were identified in progressive than regressive lesions (p=0.077; Extended Data Figure 4a-b). This remained true when the analysis was limited to clonal neoantigens (p=0.034) and there was no difference in the proportion of neoantigens that were clonal (p=0.24) (Extended Data Figure 4c-d). Further, the ratio of observed to expected neoantigens was not different (p=0.94) and there were no significant differences in binding affinity (p=0.45) or differential agretopicity index (p=0.58; Extended Data Figure 4e-h), therefore the putative neoantigens themselves were not qualitatively different in the regressive group. The increased number of neoantigens identified in progressive lesions suggests that immune escape mechanisms must be active in these lesions; indeed, these antigens may act as a selection pressure to promote the development of immune escape(9). Importantly, no overlap in tumor neoantigens was observed between different patients suggesting that vaccine-based approaches aiming to prevent progression will most likely need to be designed on a personalised basis.

Given that neoantigens are present in progressive lesions, we assessed the ability of these lesions to present antigens to the immune system. Genomic, epigenetic and transcriptomic aberrations in genes involved in MHC Class I antigen presentation (Table S3) were more prevalent in progressive than regressive lesions (p=3.9×10^−6^; Figure 3; Table S4). Considering only genomic aberrations, these were more prevalent in progressive lesions (p=0.0009) and this remained true after correcting for overall mutational burden (p=0.01), suggesting that these mutations may be under positive selection. At least one genomic aberration in MHC-associated genes was found in 25/29 progressive lesions (86%) and 5/10 regressive lesions (50%); progressive lesions had a median of 6 such changes whereas regressive lesions had a median of 0.5. Loss of heterozygosity (LOH) in the HLA region, which is found in 61% of LUSC patients(10), was identified in 34% of patients with CIS lesions. Interestingly, a similar proportion of LUSC patients (28%) demonstrated *clonal* HLA LOH, suggesting that such clonal events occur before tumor invasion. We did not find a statistically significant difference in the prevalence of HLA LOH between progressive and regressive lesions (p=0.43) although numbers were small. Expression of *HLA-A* was reduced in progressive compared to regressive lesions (p=1.9×10^−10^).

**Fig. 3.**
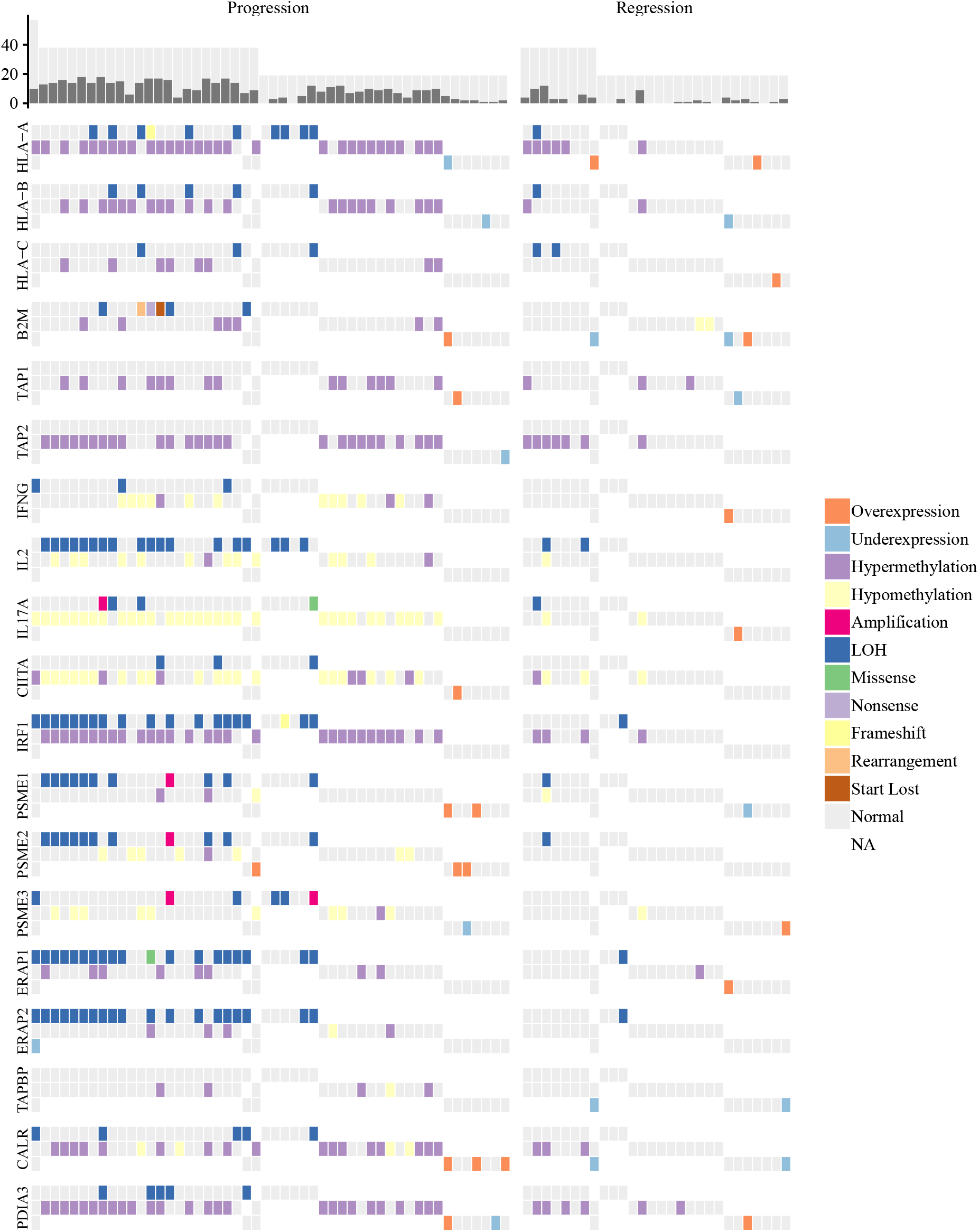
Genomic, epigenetic and transcriptomic aberrations affecting antigen-presenting genes in lung carcinoma in-situ lesions. All samples are shown (n=78; 50 progressive, 28 regressive). For each gene involved in the MHC class I pathway, aberrations are shown in transcriptomic, epigenetic and genomic data in the top, middle and bottom rows, respectively. Three genes without any identified aberrations are excluded (*CNX, HSPA, HSPC*). Samples without data for a particular modality are marked in white. The bar chart shows the number of aberrations in these genes (orange) as a proportion of the total number of possible aberrations (grey), based on the number of profiling modalities performed on each sample. Transcriptomic over/underexpression is defined as a z-score greater than ±2. Similarly, for methylation, hyper/hypomethylation is defined as z-score calculated for mean methylation beta value across the gene greater than ±2. All samples with a genomic aberration passing filters are highlighted; low-impact mutations are excluded. LOH calls integrate data from ASCAT and LOHHLA. Using a mixed-effects model to account for samples from the same patient, aberrations in this pathway are more common in progressive than regressive lesions (p=3.9×10^−6^). Considering only genomic aberrations, these were more prevalent in progressive lesions (p=0.0009) and this remained true after correcting for overall mutational burden (p=0.01).

Additionally, hypermethylation of the HLA region, which is well-described in invasive cancers(11, 12), was commonly observed, suggesting that epigenetic HLA silencing may be an important immune escape mechanism in pre-invasive disease. Genome-wide methylation analysis identified differentially methylated regions (DMRs) including a striking cluster of hypermethylation in chromosome 6 ((8); Extended Data Figure 5), covering a region containing all of the major HLA genes. This cluster was also identified in analysis of 370 LUSC versus 42 control samples published by the Cancer Genome Atlas(13). Further analysis of TCGA data demonstrate strong evidence for epigenetic silencing of multiple genes in the antigen presentation pathway: mean methylation beta value over the gene is inversely correlated with expression for *HLA-A* (r^2^=−0.32, p=2.5×10^−10^), *HLA-B* (r^2^=−0.42, <2.2×10^−16^), *HLA-C* (r^2^=−0.18, p=3.6×10^−4^), *TAP1* (r^2^=−0.53, <2.2×10^−16^) and *B2M* (r^2^=−0.38, p=1.1×10^−14^). Similar trends were observed in CIS data (Extended Data Figure 6). The methylation pattern affecting these genes is predominantly promoter hypermethylation (Extended Data Figure 7).

Demethylating agents have been shown to promote immune activation through improved antigen presentation, immune migration and T cell activity(14–16). These data support the case for moving on-going trials of demethylating agents in combination with immunotherapy from advanced lung cancer(17, 18) into early disease. Additionally, several other cancer-associated pathways are known to be affected by methylation changes(8), therefore the benefits of these drugs may extend beyond immune activation. Nevertheless, we note with caution that some key immune genes demonstrate positive correlations in TCGA data between gene expression and methylation, including the immune co-stimulating ligand *TNFSF9* (coding for 4-1BBL) (r^2^=0.32, p=1.7×10^−10^) and the MHC class II transcriptional activator *CIITA* (r^2^=0.39, p=2.5×10^−15^) (Extended Data Figure 6). Further studies will be required to demonstrate that immunological benefits of demethylating agents are not outweighed by effects on these important pathways.

Despite this evidence for impairment of antigen presentation mechanisms in CIS, we do observe ‘immune hot’ CIS lesions which progress to cancer. Next, we considered functional and microenvironment-related mechanisms to explain how these lesions were able to evade immune predation.

To study microenvironment effects on the immune response, we performed gene expression profiling on laser-captured stromal tissue taken from regions adjacent to CIS lesions. In contrast to data from gastrointestinal pre-invasive lesions(19), no genes were significantly differentially expressed on comparing stromal expression between progressive (n=10) and regressive (n=8) lesions when a FDR of <0.1 was applied. This result holds true with restricted hypothesis testing considering only genes that are related to immunity and inflammation (Figure 4a-b; Table S3).

Recent studies have identified TGF-beta signaling as a cause of T cell exclusion from tumors(20, 21), and as a potential therapeutic target(22). Whilst TGF-beta is variably expressed between progressive and regressive samples, the common downstream mediator *SMAD4* is upregulated in progressive lesions, both in CIS tissue (p=0.023) and adjacent stroma (p=0.003; Figure 4c), potentially indicating increased TGFbeta signaling in progressive lesions. Supportive of this concept, we also observed an inverse correlation between increased stromal expression of a published fibroblast TGF-beta response (FTGFB) signature(22) and TIL gradient, defined here as (TIL score in tissue) – (TIL score in stroma) (r^2^=−0.66; p=0.0029; Figure 4d). We therefore propose TGF-beta driven T cell sequestration as an additional immune escape mechanism in a subset of progressive cases. Additionally, we found upregulation of epithelial-mesenchymal transition (EMT)-related genes(23), specifically those annotated as oncogenes or with dual oncogene/tumor suppressor roles in progressive samples (Figure 4e). EMT gene expression correlated with the FTGFB signature (Figure 4f), suggesting that the immune evasion role of TGF-beta may be mediated via dysfunctional EMT transcriptional signaling affecting the tumor microenvironment, as has been previously suggested(24).

**Fig. 4.**
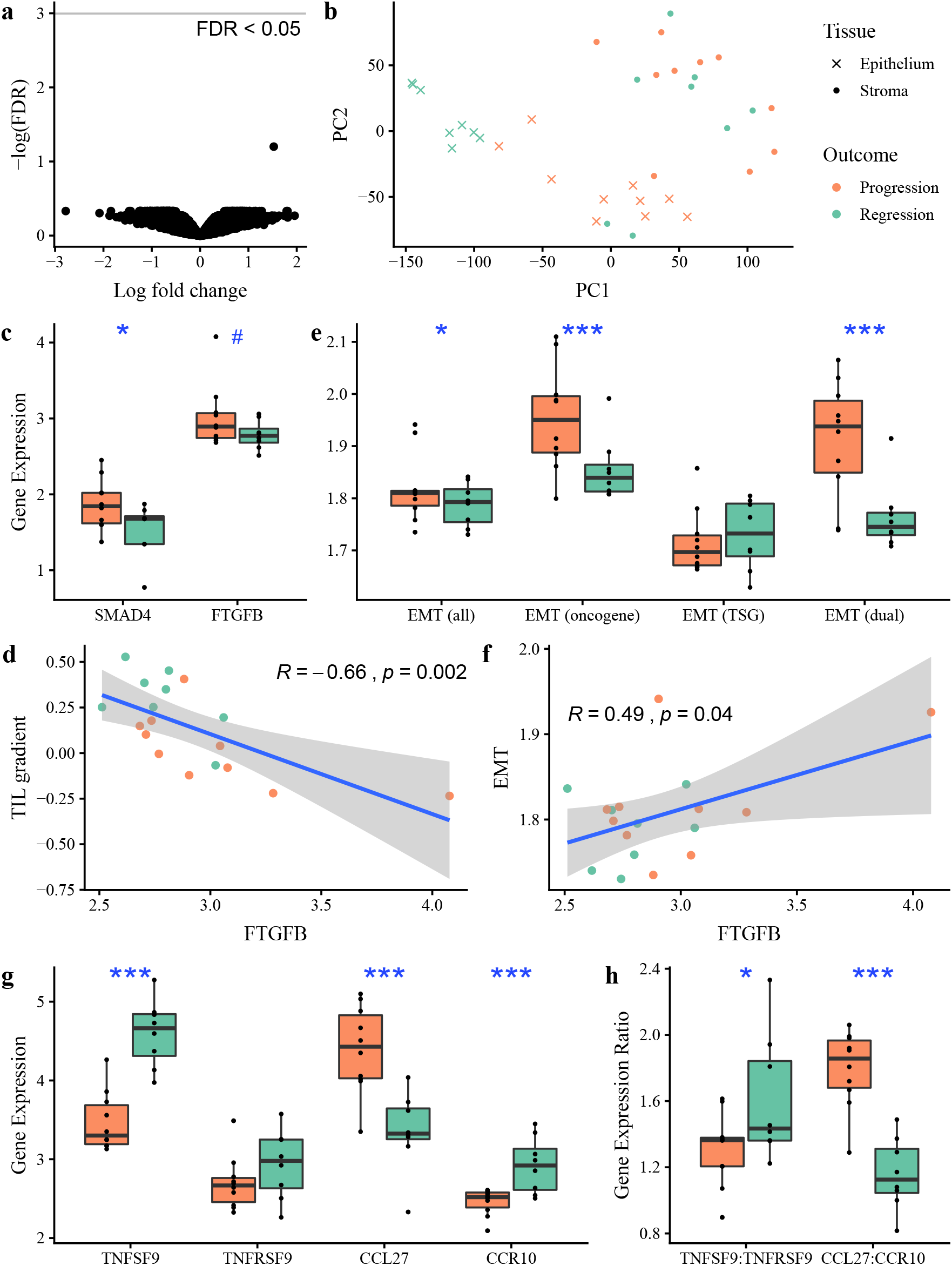
Immune escape mechanisms in CIS beyond antigen presentation. (a) Volcano plot of gene expression differential analysis of laser-captured stroma comparing progressive (n=10) and regressive (n=8) CIS samples. No genes were significant with FDR < 0.05 following adjustment for multiple testing. (b) Principle Component Analysis plot of the same 18 CIS samples, showing laser-captured epithelium and matched stroma. (c) TGF-beta signaling is increased in progressive samples, as evidenced by increased expression of the downstream gene *SMAD4* (p=0.02) and of a fibroblast TGF-beta response (FTGFBR) signature measured in matched stroma (p=0.05). The FTGFB signature, as a proxy for TGF-beta signaling, correlates inversely with TIL gradient, defined as tissue TIL score – stromal TIL score (d; r=−0.66, p=0.003). (e) EMT genes are upregulated in progressive and regressive samples. Specifically, we see upregulation of genes annotated as oncogenes (p=2.4×10^−5^) and dual oncogene/tumour suppressor functions (p=2.6×10^−5^) but not tumour suppressor genes (p=0.62). In each case we compare the geometric mean of genes in a published gene set for each sample. (f) Expression of EMT genes correlates well with the FTGFB signature (r=0.49, p=0.04). (g-h) On differential analysis of 28 immunomodulatory molecules, only *TNFSF9* was significantly upregulated (FDR 4.3×10^−5^). There was no corresponding upregulation of the *TNFRSF9* receptor. A comparison of ligand:receptor ratios for known cytokines identified only *CCL27:CCR10* as upregulated in progressive samples (FDR 0.003). All p-values are calculated using linear mixed effects modeling to account for samples from the same patient; ***p < 0.001 **p < 0.01 *<0.05 ^#^<0.1. Units for gene expression figures represent normalised microarray intensity values.

To identify differences in cytokine responses between progressive and regressive lesions, we calculated the ligand:receptor mRNA expression ratio for 52 known cytokine:receptor pairs(25). Only one, *CCL27:CCR10*, was significant with FDR < 0.01 (Fold change 1.55, FDR 0.003); progressive samples express more *CCL27* (p=2.6×10^−6^) and less *CCR10* (p=0.1×10^−4^) than regressive (Figure 4g-h). *CCL27:CCR10* signaling has been associated with immune escape in melanoma through PIK/Akt activation in a mouse model(26); in CIS, *CCL27* expression correlates with expression of both *PIK3CA* (r^2^=0.61, p=0.008) and *AKT1* (r^2^=0.68, p=0.002) (Extended Data Figure 8). *CCL27* is minimally expressed in both normal lung tissue and invasive squamous cell lung cancer(13, 27), suggesting that this effect is specific to early carcinogenesis and therefore warrants further investigation as a target for preventative therapy.

Targeting immunomodulatory molecules such as PD-1 now forms part of first-line lung cancer management(28). To investigate the role of such molecules in pre-invasive immune escape, we performed differential expression analysis between progressive and regressive lesions, focused on 28 known immunomodulatory genes (Table S3). *TNFSF9* (4-1BBL, CD137L) was significantly downregulated in progressive lesions (FDR=4.34×10^−5^; Figure 4g-h) with no corresponding change identified in its receptor *TNFRSF9* (FDR=0.6). *TNFSF9* promotes activation of T cells and natural killer (NK) cells(29); in CIS lesions *TNFSF9* expression correlates with cytotoxic cell (r^2^=0.77, p=0.0002) and NK cell infiltration (r^2^=0.54, p=0.02), as predicted from gene expression data. Agonists of the *TNFSF9* receptor have been shown to be clinically efficacious in several cancers(30–32) and these data support their investigation in targeted early lung cancer cohorts. Furthermore, individual lesions showed notably high or low expression of other immunomodulatory genes, raising the possibility that other immunomodulators may be targets for therapy in individual cases (Extended Data Figure 9).

Our previous work highlighted occasional cases of ‘late progressive’ lesions, which met a clinical endpoint of regression (defined by the subsequent biopsy at the same site showing resolution to normal epithelium or low-grade dysplasia) but the index CIS biopsy had the molecular appearance of a progressive lesion, and it indeed subsequently developed cancer months or years later. Clinical review identified 11 lesions across the 53 regressive lesions in our current cohort (20.7%) that at later clinical follow up subsequently progressed to cancer, and hence are termed ‘late progressive’. These included 4 previously published lesions subjected to whole-genome sequencing and/or methylation and shown to display the genomically unstable appearance of progressive lesions, as well as 7 with immunohistochemistry data and 10 with lymphocyte quantification performed from H&E slides (Table S1; Extended Data Figure 1). Interestingly, based on these data, late progressive lesions appear immunologically similar to regressive lesions, showing increased infiltration with lymphocytes and CD8 cells compared to progressive lesions (Extended Data Figure 10).

Whilst we acknowledge that sample numbers are small when examining subgroups of regressive lesions in this way, our data support a model in which lesions should be considered on two axes: genomic stability and immune competence. Our previous work predicts that chromosomally unstable lesions will usually progress, implying that they have escaped immune predation. Yet some may regress if they remain immune competent only to later progress, potentially due to their genomic instability making them more likely to evolve immune escape mechanisms during regression, and hence become ‘late progressors’. Of 11 late progressors in this cohort, median time from regressive index biopsy to progression was 3.2 years (range 0.8-4.6 years). This time period represents a change from a point of known immune competence to demonstrated immune escape. Hence, we might estimate that a successful therapeutic strategy to block a particular immune escape mechanism might delay the onset of cancer by around 3 years. Of the remaining 42 regressive samples in this cohort, median follow-up time was 4.73 years (range 0.42-13.5 years), suggesting that genomically ‘stable’ samples are likely to regress and remain regressed long-term. Given their immunological competence, late progressors are included in the regressive cohort when analysing immune escape mechanisms in this study.

In summary, we present evidence that immune surveillance may play a critical role in spontaneous regression of pre-cancerous lesions of the airways. We identify mechanisms of immune escape present before the point of cancer invasion, many of which offer potential therapeutic targets. Analysis of ‘late progressive’ samples provides insight into the dynamics of this process. These data present an opportunity to induce regression and prevent cancer development. Demethylating agents, 4-1BB agonists, CCL27 and TGF-beta blockade are therapeutic candidates that warrant further research. As a result of field carcinogenesis, patients with pre-invasive lesions are at risk of synchronous cancers at other sites, which are likely to be clonally related(8, 33) and therefore may benefit from systemic immunomodulatory treatment. The data presented here support a new paradigm of personalised immune-based systemic therapy in early disease.

## Methods

### Ethical approval

All tissue and bronchial brushing samples were obtained under written informed patient consent and were fully anonymized. Study approval was provided by the UCL/UCLH Local Ethics Committee (REC references 06/Q0505/12 and 01/0148). All relevant ethical regulations were followed.

### Cohort description and patient characteristics

For over 20 years, patients presenting with pre-invasive lesions, which are precursors of squamous cell lung cancer (LUSC), have been referred to the UCLH Surveillance Study. As previously described(1), patients undergo repeat bronchoscopy every four months, with definitive treatment performed only on detection of invasive cancer. Autofluorescence bronchoscopy is used to ensure the same anatomical site is biopsied at each time point. Gene expression, methylation and whole genome sequencing data of carcinoma in-situ (CIS) samples have been performed on this cohort, and data have been published(2). These data are used in this study.

All patients enrolled in the UCLH Surveillance Study who met a clinical end point of progression or regression were included; by definition they underwent an ‘index’ CIS biopsy followed by a diagnostic cancer biopsy (progression) or a normal/low-grade biopsy (regression) four months later. Index lesions were identified between 1999 and 2017. Cases meeting an end-point of regression underwent clinical review to identify those which subsequently progressed; 11 samples (20.7%) were identified, which are described as ‘late progressors’ in the main text. Of these 11, median time from ‘regressive’ index biopsy to progression was 3.2 years (range 0.8-4.6 years) whilst the remaining 42 samples had a median follow up time of 4.73 years (range 0.42-13.5 years). Whilst we cannot fully exclude that any regressive sample may later develop cancer, the fact that median follow up in the study group was longer than the maximum follow up in the late progression group suggests that late progression in included samples is unlikely.

All samples underwent laser capture microdissection (LCM) to ensure only CIS cells underwent molecular profiling. Methods for sample acquisition, quality control and mutation calling are as previously described, as are full details regarding patient clinical characteristics.

Briefly, gene expression profiling was performed using both Illumina and Affymetrix microarray platforms. Normalisation was performed using proprietary Illumina software and the RMA method of the affy(3) Bioconductor package respectively. This study includes 18 previously unpublished gene expression arrays from stromal tissue. These samples were collected using LCM to identify stromal regions adjacent to 18 already-published CIS samples (corresponding to the 18 samples undergoing Affymetrix microarray profiling described above). These new stromal samples underwent Affymetrix profiling using the exact same methodology as previously described for CIS tissue samples. To avoid issues related to batch effects between platforms, the analyses in this paper utilise only samples profiled on Affymetrix microarrays, which include both CIS and matched stromal samples. Methylation profiling was performed using the Illumina HumanMethylation450k microarray platform. All data processing was performed using the ChAMP Bioconductor package(4).

For both gene expression and methylation data, z-scores were used to identify significant aberrations. These were calculated using regressive samples as a reference cohort for gene expression data, and control brushings for methylation data. Whole genome sequencing data was obtained using the Illumina HiSeq X Ten system. A minimum sequencing depth of 40x was required. BWA-MEM was used to align data to the human genome (NCBI build 37). Unmapped reads and PCR duplicates were remoted. Substitutions, insertions-deletions, copy number aberrations and structural rearrangements were called using CaVEMan(5), Pindel(6, 7), ASCAT(8) and Brass(9) respectively.

### Comparison of Microarray Platforms

As described above, our previous work performed gene expression profiling using both Illumina and Affymetrix microarray platforms (GEO platform IDs GPL13534 and GPL18281 respectively), with Illumina data used for discovery analysis and Affymetrix as a validation set. Our previous publication did not identify clear differences in immune pathways between progressive and regressive lesions based on the Illumina discovery set, yet a similar analysis of the Affymetrix dataset does identify two significant immune-related KEGG pathways(10): cytokine-cytokine interaction (hsa04060) and type I diabetes mellitus (hsa04940). We therefore questioned whether this disparity may be due to platform differences. The Affymetrix platform used has many more probes than the Illumina platform, allowing coverage of more genes and coverage of multiple transcripts for some genes. To examine the impact of these differences we performed pathway analysis on the Illumina and Affymetrix datasets separately, then repeated this analysis using only probes that were shared by both platforms and were unambiguous (i.e. had a one-to-one mapping to a given gene on both microarray platforms). Using a Gene Set Enrichment Analysis (GSEA) method, we found two immune-related KEGG pathways to be significant in the Affymetrix dataset but not the Illumina dataset: cytokine-cytokine interaction (hsa04060) and type I diabetes mellitus (hsa04940). Both of these pathways included genes which were not profiled in the Illumina dataset, and indeed when the Affymetrix dataset was reduced to include only shared unambiguous probes hsa04940 was no longer significant and hsa04060 showed a smaller effect size. Chromosomal instability related genes – the most important finding from our previous work – remained significant across all analyses. Some genes which are important to our present analysis are not covered by the Illumina microarray, including *TNFSF9*, *CXCL8* and *CD274*. We believe these differences justify our decision to focus on the Affymetrix platform, as it offers wider coverage of important immune genes. Pathway analysis results are included in Supplementary Table 5.

### Sample selection for profiling

As previously described, all patients enrolled in the surveillance programme discussed above were considered for this study. For a given CIS lesion under surveillance, when a biopsy from the same site in the lung showed evidence of progression to invasive cancer or regression to normal epithelium or low-grade dysplasia, we defined the preceding CIS biopsy as a progressive or regressive ‘index’ lesion respectively. Due to the small size of bronchoscopic biopsy samples, not all profiling techniques were applied to all samples. Patients with Fresh Frozen (FF) samples underwent whole genome sequencing and/or methylation analysis depending on sample quality. Patients with formalin-fixed paraffin-embedded (FFPE) samples underwent gene expression analysis. Further detail is available in our previous manuscript(2). Additionally, any patient with an available FFPE block underwent image analysis as described below, and all patients with Affymetrixbased gene expression profiling underwent further profiling of laser-captured adjacent stroma.

### Statistical Methods

Unless otherwise specified, all analyses were performed in an R statistical environment (v3.5.0; www.r-project.org/) using Bioconductor(11) version 3.7. Code to reproduce a specific statistical test is publicly available at the Github repository below.

Unless otherwise stated, comparisons of means between two independent groups are performed using a two-sided Wilcoxon test. In some cases, multiple samples have been profiled from the same patient, although always from distinct sites within the lung. In such cases we used mixed effects models to compare means between groups, treating the patient ID as a random effect, as implemented in the Bioconductor *lme4* library(12), with p-values calculated using the Anova method from the Bioconductor *car* library(13). Differential expression was performed using the *limma*(14) Bioconductor package to compare microarray data between two groups. When adjustment for multiple correction is required we quote a False Discovery Rate (FDR) which is calculated using the Benjamini-Hochberg method(15). Cluster analysis and visualization was performed using the *pheatmap*(16) Bioconductor package.

### Image analysis

All slides were scanned using NanoZoomer Digital Pathology System scanner model C9600-01, using NDP.scan version 2.5.89 (Hamamatsu, Japan).

Four distinct cell types from H&E images were identified with an automated deep learning pipeline trained using 21,009 pathological annotations from NSCLC samples in the TRACERx100 cohort(17). The four classes correspond to cancer cells, lymphocytes that included leukocytes and plasma cells, stromal cells that included fibroblasts and endothelial cells, and an “other” cell type that included nonidentifiable and less abundant cells such as macrophages, chondrocytes, and pneumocytes. Customised implementation of spatially constrained convolution neural networks(18) for TensorFlow were used for the single cell classification and detection tasks. The deep learning pipeline was validated using 5,951 pathological annotations within TRACERx as well as 5,082 annotations collected externally on an independent cohort of 100 NSCLC cases from the LATTICe-A study(19). Biological validation of this algorithm against immunohistochemistry data has been previously described (submitted for publication).

### IHC

2-5μm tissue sections were cut and transferred onto poly-l-lysine–coated slides, dewaxed in two changes of xylene and rehydrated in a series of graded alcohols. Details of the three primary antibodies used are as follows:

- SP35: Anti-CD4 Rabbit monoclonal antibody from Spring Biosciences Inc., Pleasanton, CA, US.
- SP239: Anti-CD8 Rabbit monoclonal antibody from Spring Biosciences Inc., Pleasanton, CA, US.
- 236A/E7: Anti-FOXP3 Mouse antibody, Kind gift from Dr G Roncador, CNIO, Madrid (Spain).

Single immunohistochemistry was carried out using the automated platforms BenchMark Ultra (Ventana/Roche) and the Bond-III Autostainer (Leica Microsystems) according to a protocol described elsewhere(20, 21). To establish optimal staining conditions (i.e. antibody dilution and incubation time, antigen retrieval protocols, suitable chromogen) each antibody was tested and optimized on sections of human reactive tonsil, used as positive control.

Multiplex immunohistochemistry was carried out using a protocol described previously(21). Co-expression of nuclear and cytoplasmic or membranous proteins was easy to detect, as the colour of the chromogens remained distinct. Specificity of the staining was assessed by a haematopathologist (TM) with expertise in multiplex-immunostaining. Slides were scanned using the Hamamatsu Nanozoomer digital scanner as described above.

For T cell subset quantification, a similar deep learning pipeline was used. The convolutional neural networks were trained on sample TRACERx IHC CD4/CD8/FOXP3 images using 9,333 pathological annotations and validated against 6 NSCLC independent images using 5,028 pathological annotations. The IHC algorithm classified cells into four classes: CD8+, CD4+, FOXP3+ and “other” cell class (hematoxylin cells). When comparing cell counts between samples, absolute counts were divided by the region area. Regions of CIS and stroma within a slide were quantified separately, with regions annotated manually by the investigators.

### Neoantigen prediction and LOHHLA

HLA typing was performed using Optitype(22) on germline (blood) WGS data from each patient. This was used as input for netMHCpan 4.0(23, 24) for neoantigen prediction; 9-, 10- and 11-mer peptides were considered for each somatic mutation, called using methods described above. To assess for quantitative differences between neoantigens in the progressive and regressive groups, we compared their binding affinities (as calculated by netMHCpan) and their differential agretopicity index (DAI), defined as the difference in binding affinity between mutant and wild-type peptides. Significant differences in these values were not observed between the regressive and progressive groups.

The same HLA typing data was used as input to the LOHHLA tool(25) (Loss of Heterozygosity in Human Leukocyte Antigen), alongside copy number, purity and ploidy data derived from ASCAT. This tool assesses each sample for the presence of LOH in the HLA region – a difficult task due to polymorphism in this region. Output plots from LOHHLA were visually checked prior to calling the presence or absence of HLA LOH in a sample.

### DMR analysis

Methylation data analysis was performed using the Chip Analysis Methylation Pipeline (*ChAMP*) Bioconductor package with default settings(4). The functions *champ.DMP()* and *champ.DMR()* were used to identify differentially methylated probes (DMPs) and differentially methylated regions (DMRs) respectively. Annotation of DMPs and DMRs with affected genes is performed by default within these functions.

A criticism raised against this analysis is the identification of DMRs affecting a highly polymorphic region of chromosome
6. However, we argue that this is a differential analysis between two groups (progressive and regressive), with results replicated in an independent dataset from TCGA (Cancer vs Control data), therefore should not be affected by polymorphism unless the underlying HLA types are significantly different between the two groups. For each identified HLA type, based on 4-digit resolution, we compared the number of patients identified in the progressive and regressive groups using a Fisher’s exact test, and did not find any HLA types to be significant with p < 0.05.

### Immune cell quantification from GXN data

To estimate relative immune cell populations from gene expression data we applied the method of Danaher et al.(26) This method was chosen as it has been shown to out-perform similar methods when benchmarked against immunohistochemistry in a large analysis of early-stage invasive lung cancer(27). Briefly, for each of 15 immune cell types, a small set of genes is defined which has been shown to correlate with the presence of that cell type. For each cell type, the mean expression of its associated genes gives a ‘score’ for that cell type. If a gene is not measured by the Affymetrix microarray used, that gene is ignored.

A ‘TIL score’, estimating the overall infiltration of lymphocytes into the tissue, is calculated by taking the mean of 10 individual cell type scores (B-cells, Cytotoxic cells, Exhausted CD8, Macrophages, Neutrophils, NK CD56dim cells, NK cells, T-cells, Th1 cells, CD8 T cells). This process is encoded in the R function *do.danaher()*, which is available from the Github repository accompanying this paper.

### Immune cell quantification from methylation data

Similar immune quantification from methylation data was performed using *methylCIBERSORT*(28). Methylation data was first converted to a mixture file using the *methylCIBERSORT* R package version 0.2.0. A signature file for squamous cell lung cancer was also taken from this package; this signature was derived from TILs in squamous cell lung cancer, a very similar biological question to that of our study. These data were used as input to CIBERSORT(29) to provide relative values for each immune cell subtype included in the signature file.

## Data Availability

All raw data used in this study is publicly available. Previously published CIS gene expression and methylation data is stored on GEO under accession number GSE108124; matched stromal gene expression data is stored under accession number GSE133690. Previously published CIS whole genome sequencing data is available from the European Genome Phenome Archive (https://www.ebi.ac.uk/ega/) under accession number EGAD00001003883.

## Code Availability

All code used in our analysis will be made available at http://github.com/uclrespiratory/cis_immunology on publication. All software dependencies, full version information, and parameters used in our analysis can be found here.

## Author Contributions

A.P. and V.H.T. contributed equally to this work, as did K.A., S.E.A.R. and T.L.. A.P., V.H.T., N.M. and S.M.J. co-wrote the manuscript. S.M.J., S.A.Q., V.H.T. and A.P. conceived the study design. V.H.T., D.C. and S.A. performed stromal LCM and gene expression profiling experiments. C.P.P. performed LCM and methylation experiments. H.L-S. and P.J.C. performed genomic experiments. A.A., T.L., J.Y.H. and T.M. designed and performed IHC experiments. K.A., S.E.A.R. and Y.Y. performed cell quantification on H&E and IHC images. S.M.J., P.J.G., B.C. and R.M.T. led the bronchoscopic surveillance programme through which samples were obtained. M.F. performed histological review. P.F.D. performed pathological processing. A.P. performed bioinformatic analysis, supported by R.R. and N.M.. R.E.H., K.H.C.G., C.D., A.F., C.S., C.T., S.A.Q. and N.M. gave advice and reviewed the manuscript. S.M.J. provided overall study oversight.

## ACKNOWLEDGEMENTS

We thank all of the patients who participated in this study. We thank P. Rabbitts, A. Banerjee and C. Read for their early development of the study. The results published here are in part based on data generated by a TCGA pilot project established by the National Cancer Institute and National Human Genome Research Institute. Information about TCGA and the investigators and institutions that constitute the TCGA research network can be found at http://cancergenome.nih.gov. R.E.H., N.M., P.J.C., and S.M.J. are supported by Wellcome Trust fellowships. S.M.J. is also supported by the Rosetrees Trust, the Welton Trust, the Garfield Weston Trust, the Stoneygate Trust and UCLH Charitable Foundation. V.T., C.P., R.E.H., S.A. and S.M.J. have been funded by the Roy Castle Lung Cancer Foundation. A.P. and D.C. are funded by Wellcome Trust clinical PhD training fellowships. H.L.-S. is funded by the Wellcome Trust Sanger Institute non-clinical PhD studentship. C.T. was a CRUK Clinician Scientist. This work was partially undertaken at UCLH/UCL, who received a proportion of funding from the Department of Health’s NIHR Biomedical Research Centre’s funding scheme (S.M.J.). R.E.H., N.M., C.S., and S.M.J. are part of the CRUK Lung Cancer Centre of Excellence. C.S., and S.M.J. are supported by Stand Up to Cancer. Y.Y. acknowledges funding from Cancer Research UK Career Establishment Award, Breast Cancer, Children’s Cancer and Leukaemia Group, NIH U54 CA217376 and R01 CA185138, CDMRP Breast Cancer Research Program Award, CRUK Brain Cancer Award (TARGET-GBM), European Commission ITN, Wellcome Trust, and The Royal Marsden/ICR National Institute of Health Research Biomedical Research Centre. S.A.Q. is funded by a CRUK Senior Cancer Research Fellowship, a CRUK Biotherapeutic Program Grant, the Cancer Immunotherapy Accelerator Award (CITA-CRUK) and the Rosetrees Trust. The funders had no role in study design, data collection and analysis, decision to publish or preparation of the manuscript.

---

**Extended Data Fig 1.**
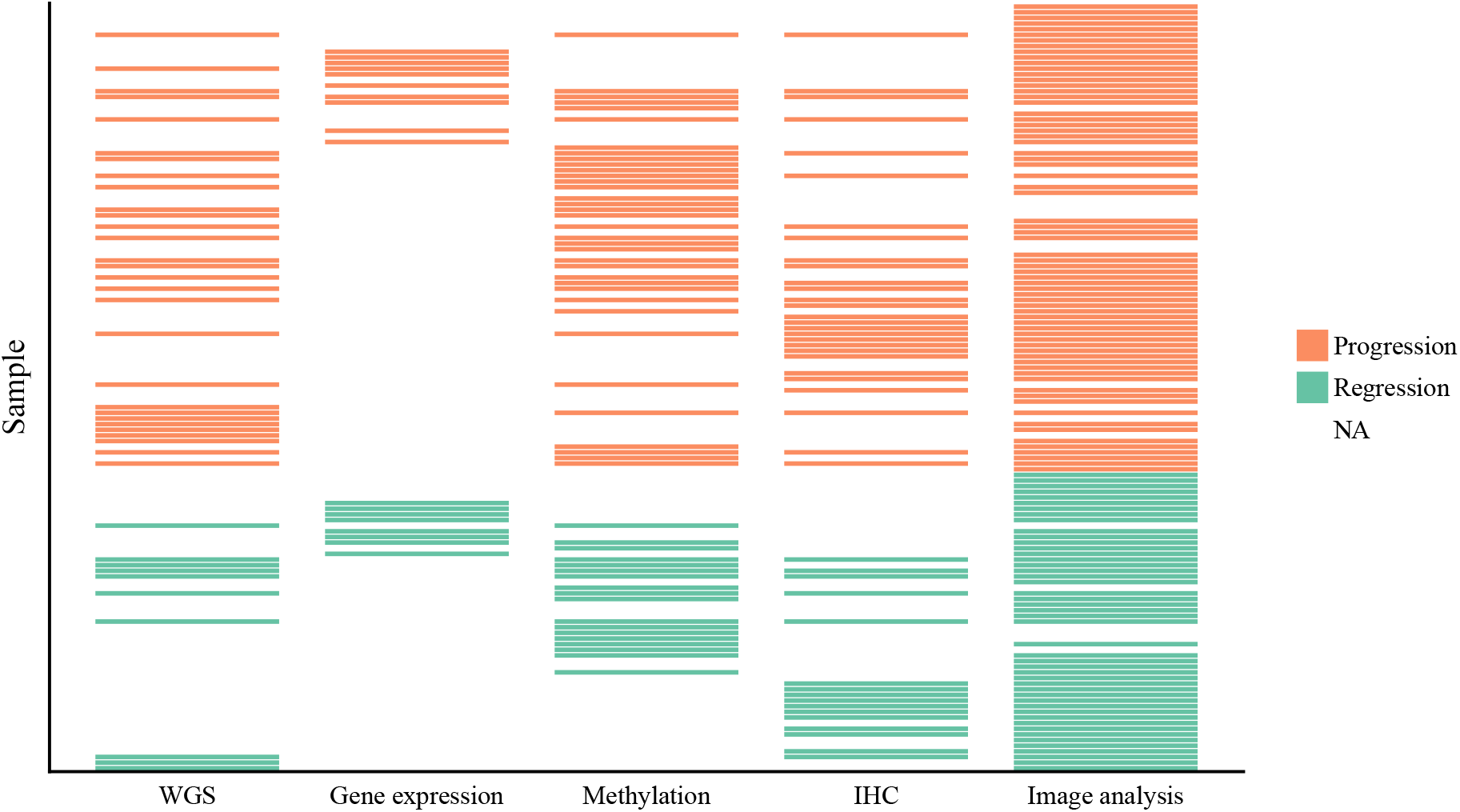
Summary of analyses performed on each CIS sample. Due to technical limitations related to the small size of bronchoscopic biopsies, not all analyses were performed on all samples. Table S1 provides a detailed reference of analyses performed on a per-sample basis. Methodology for sample selection for each analysis modality is provided in methods.

**Extended Data Fig 2.**
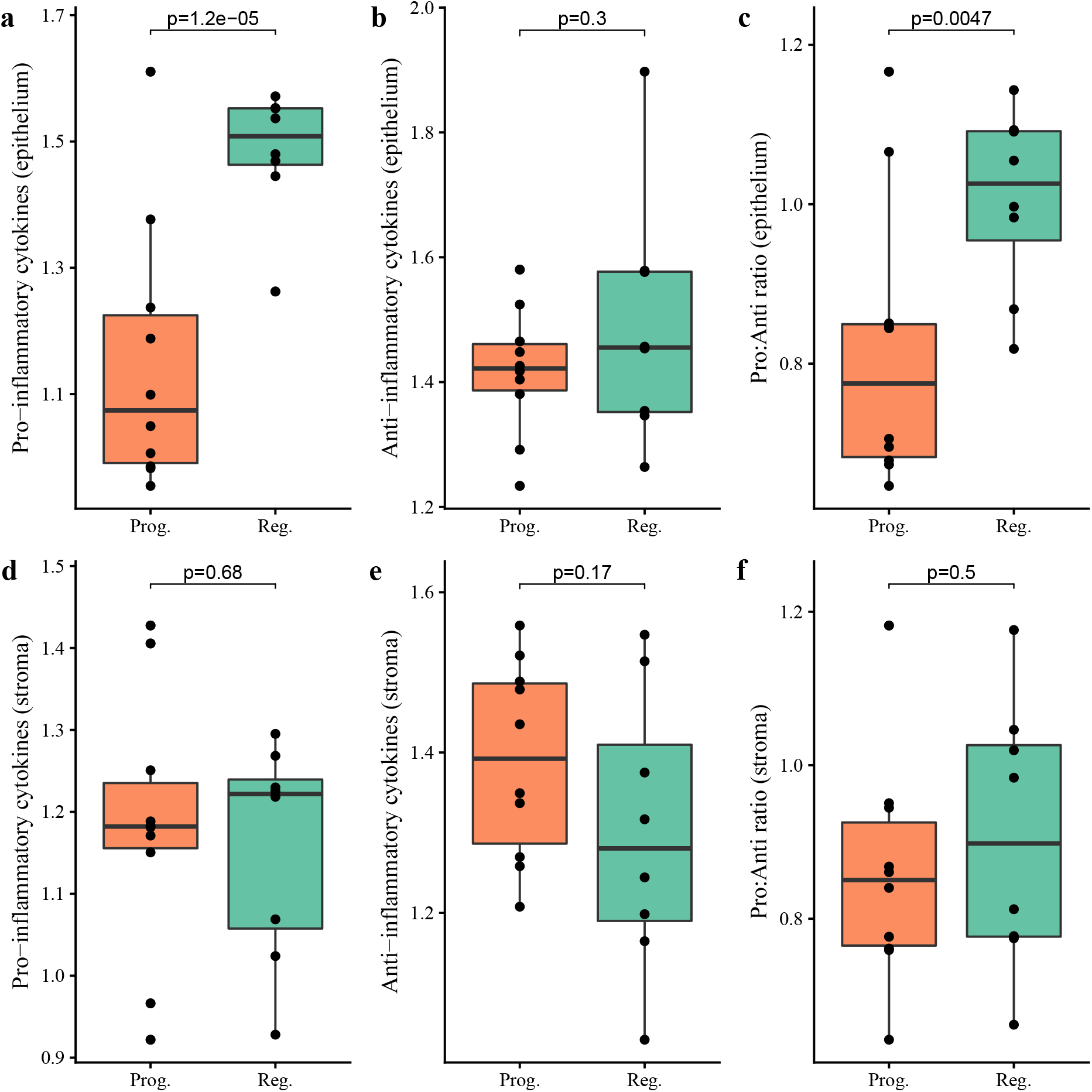
Comparing transcriptomic data from progressive CIS lesions (n=10) with regressive (n=8) we find regressive lesions express higher levels of proinflammatory cytokines (a) but not anti-inflammatory cytokines (b) within the epithelium. The pro:anti-inflammatory ratio is higher in regressive lesions (c). Transcriptomic data from laser-captured stroma adjacent to the same lesions does not show any difference in cytokine expression between progressive and regressive lesions (d-f). Expression values shown are the geometric means of gene expression data for 9 pro-inflammatory and 7 anti-inflammatory cytokines. p-values are calculated using linear mixed effects modelling to account for samples from the same patient.

**Extended Data Fig 3.**
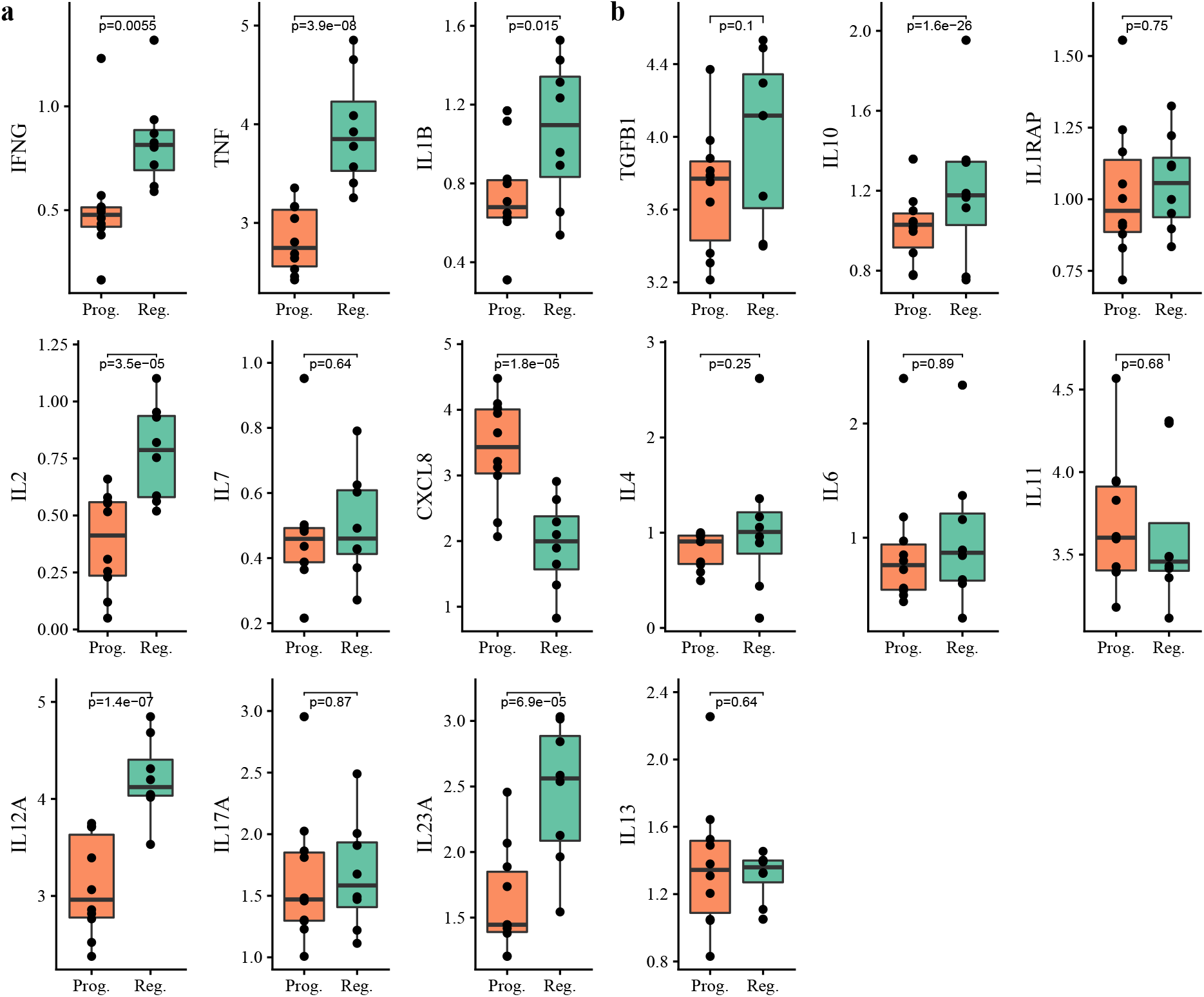
Expression of individual cytokines in progressive and regressive CIS lesions. Continuing the analysis of transcriptomic data from progressive CIS lesions (n=10) with regressive (n=8) shown in Extended Data Figure 2, we demonstrate the contributions of individual pro-inflammatory cytokines (a) and anti-inflammatory cytokines (b). We see upregulation of several pro-inflammatory cytokines in regressive lesions: *IFNG, IL12A, IL2, IL23A* and *TNF*, as well as the classically anti-inflammatory cytokine *IL10*. *CXCL8*, which is associated with macrophages, is downregulated in regressive lesions. p-values are calculated using linear mixed effects modeling to account for samples from the same patient.

**Extended Data Fig 4.**
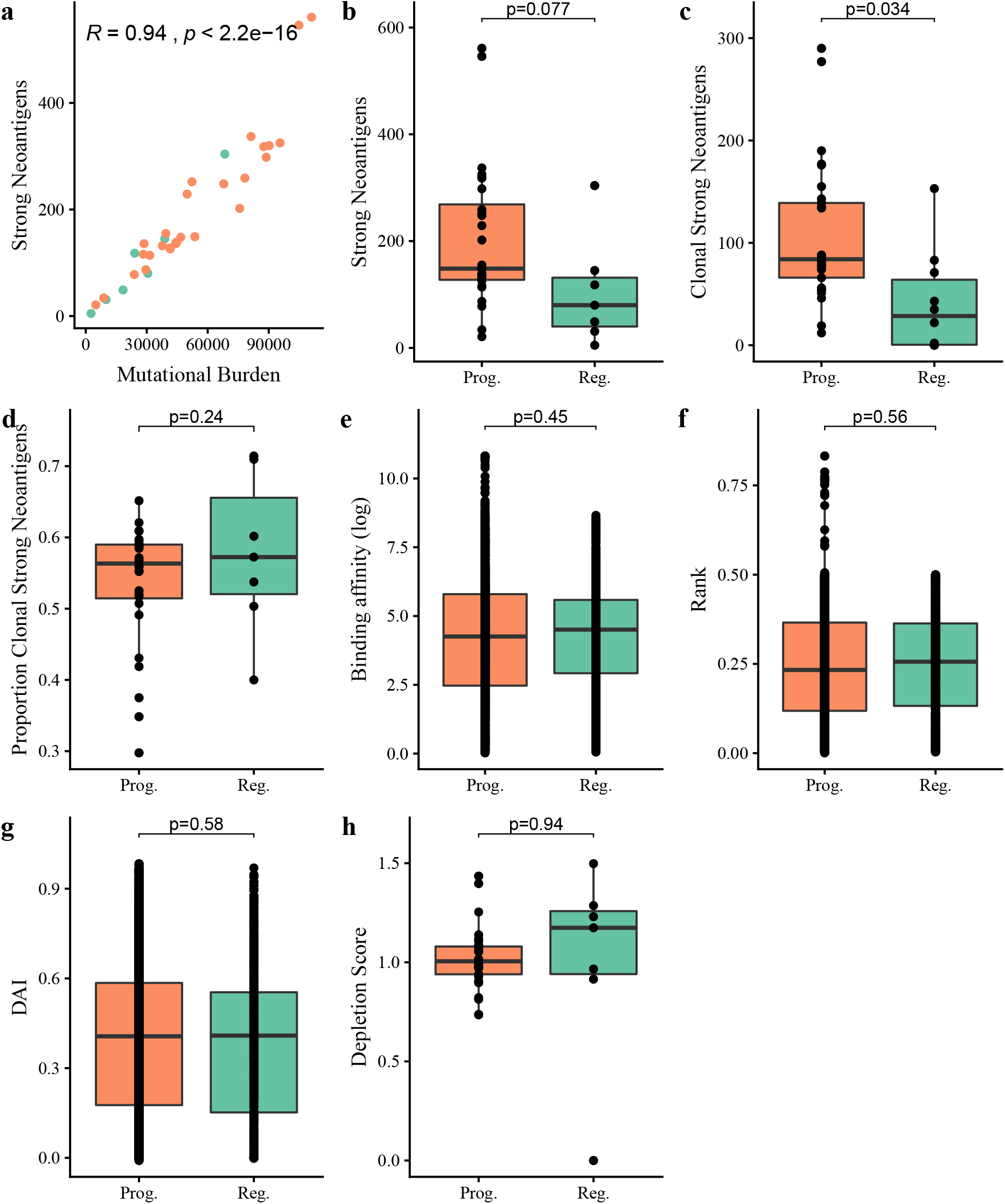
Neoantigen analysis of progressive versus regressive lesions. Predicted neoantigen load correlates closely with mutational burden (a). Therefore, progressive samples, which harbor more mutations, have more neoantigens (b). This remains true when the analysis is limited to clonal neoantigens (c). The proportion of clonal neoantigens was similar (d). Considering the individual predicted neoantigens, there was no qualitative difference between progressive and regressive samples; they were similar in terms of binding affinity (e), rank binding affinity (f) and differential agretopicity index (DAI) (g). The ratio of observed to expected neoantigens (‘depletion score’) was similar between progressive and regressive lesions (h). The p-value for figure (a) was calculated using Pearson’s product moment; p-values for figures (b)-(h) were calculated using a Wilcoxon rank-sum test.

**Extended Data Fig 5.**
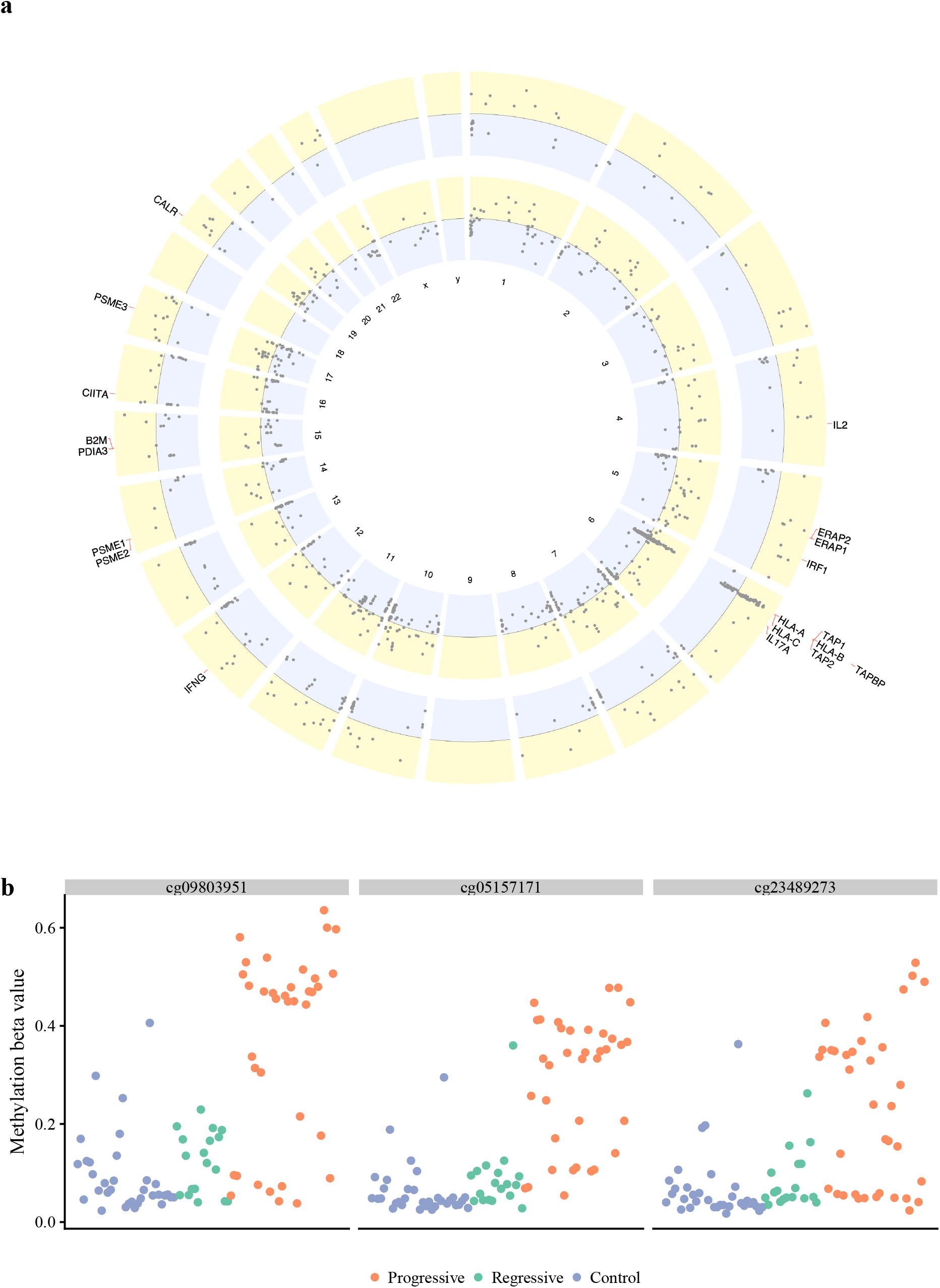
Aberrant methylation of the HLA region is a feature of progressive CIS and cancer. (a) Differentially methylated regions across the genome, calculated for progressive vs regressive CIS (outer circle) and for cancer vs control (inner circle). Hypermethylated DMRs are plotted in yellow, hypomethylated in blue. Genes involved in the MHC class I mechanism are highlighted. In both comparisons a cluster is observed on chromosome 6, which includes all main HLA regions. (b) Selection of three probes covering the *HLA-A* gene, all showing marked hypermethylation in a subset of progressive samples and hence suggesting an epigenetic mechanism for reduced *HLA-A* in these samples.

**Extended Data Fig 6.**
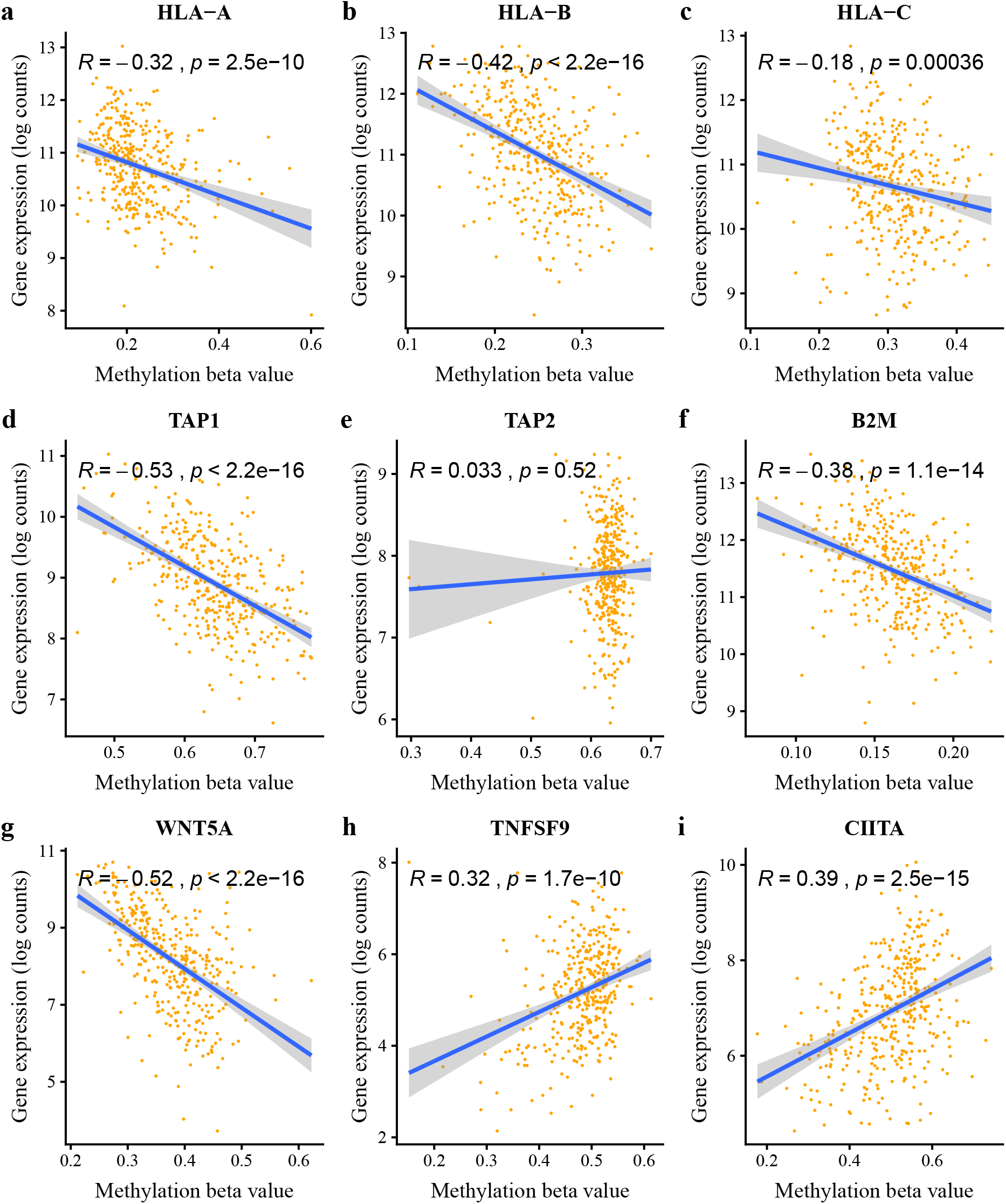
Epigenetic silencing of antigen-presenting genes in squamous cell lung cancers. (a-f) Correlations of expression and methylation data from TCGA for key antigen-presentation genes demonstrates clear evidence of epigenetic silencing. Silencing is also seen for other cancer-associated genes such as *WNT5A* (g), suggesting that demethylating agents may have wider benefits than improving antigen presentation. However, some key immune genes including immunomodulatory molecule *TNFSF9* (h) and MHC II regulator *CIITA* (i) show a positive correlation with methylation, suggesting that demethylating agents may not be universally beneficial on the immune response. Correlation coefficients shown are calculated using Pearson’s product moment.

**Extended Data Fig 7.**
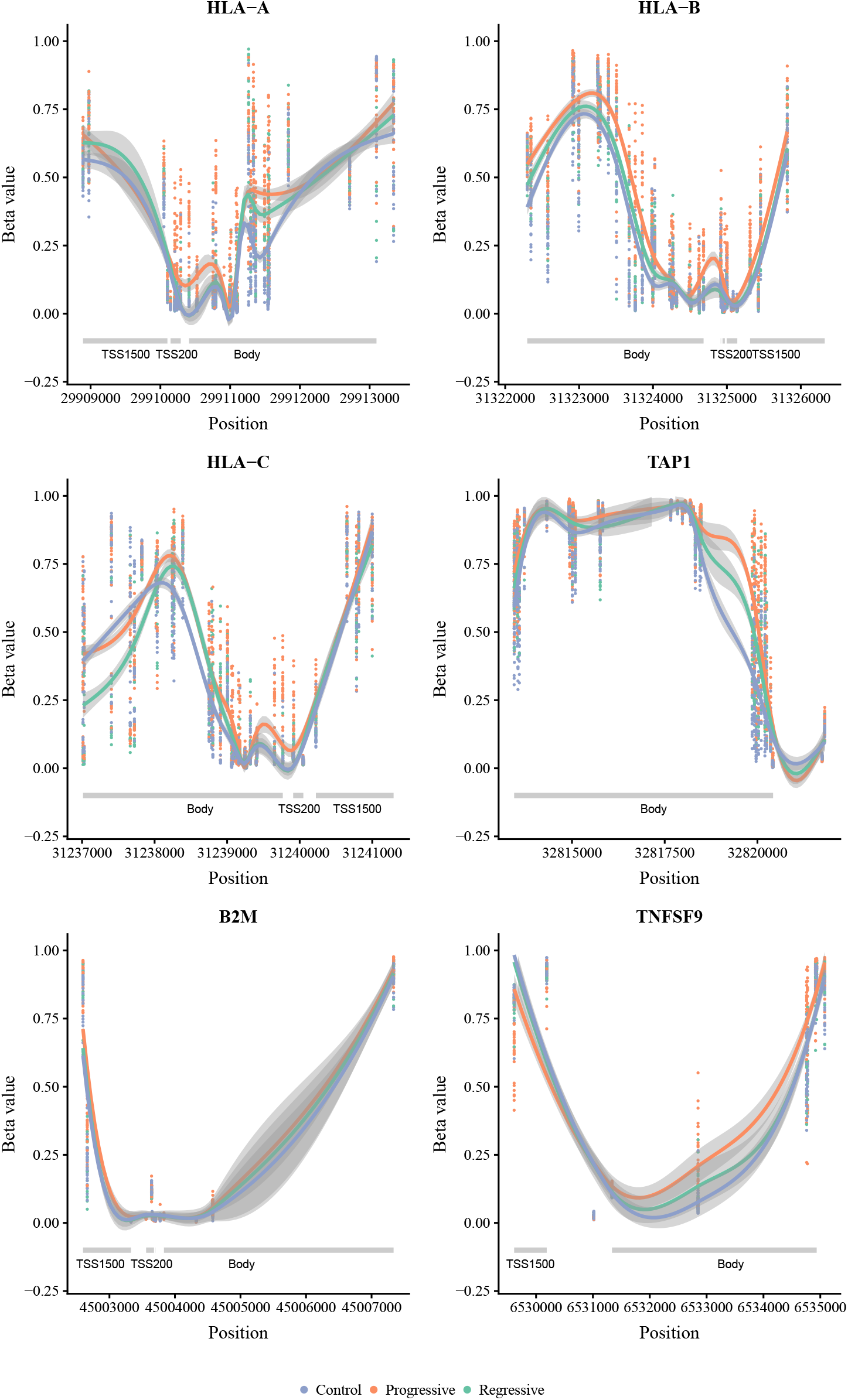
Methylation patterns over antigen-presenting genes. Methylation patterns are shown for antigen presentation genes *HLA-A, HLA-B, HLA-C, TAP1* and *B2M*, as well as the immunomodulator *TNFSF9*. Methylation data is generated from Illumina 450k microarrays, which measure methylation at 450,000 probes across the genome. In each plot, the x-axis shows the genomic location of each probe related to the gene of interest. On the y-axis, probe values are shown for each sample, coloured as progressive (red; n=36), regressive (green; n=18) or control (blue; n=33). Loess lines for each sample group are shown, with error bars in grey. We see a pattern of promoter hypermethylation in progressive samples for the majority of these genes, consistent with epigenetic silencing. An exception is *TNFSF9* which shows predominantly body hypermethylation; this is consistent with the observation that hypermethylation of *TNFSF9* increases expression.

**Extended Data Fig 8.**
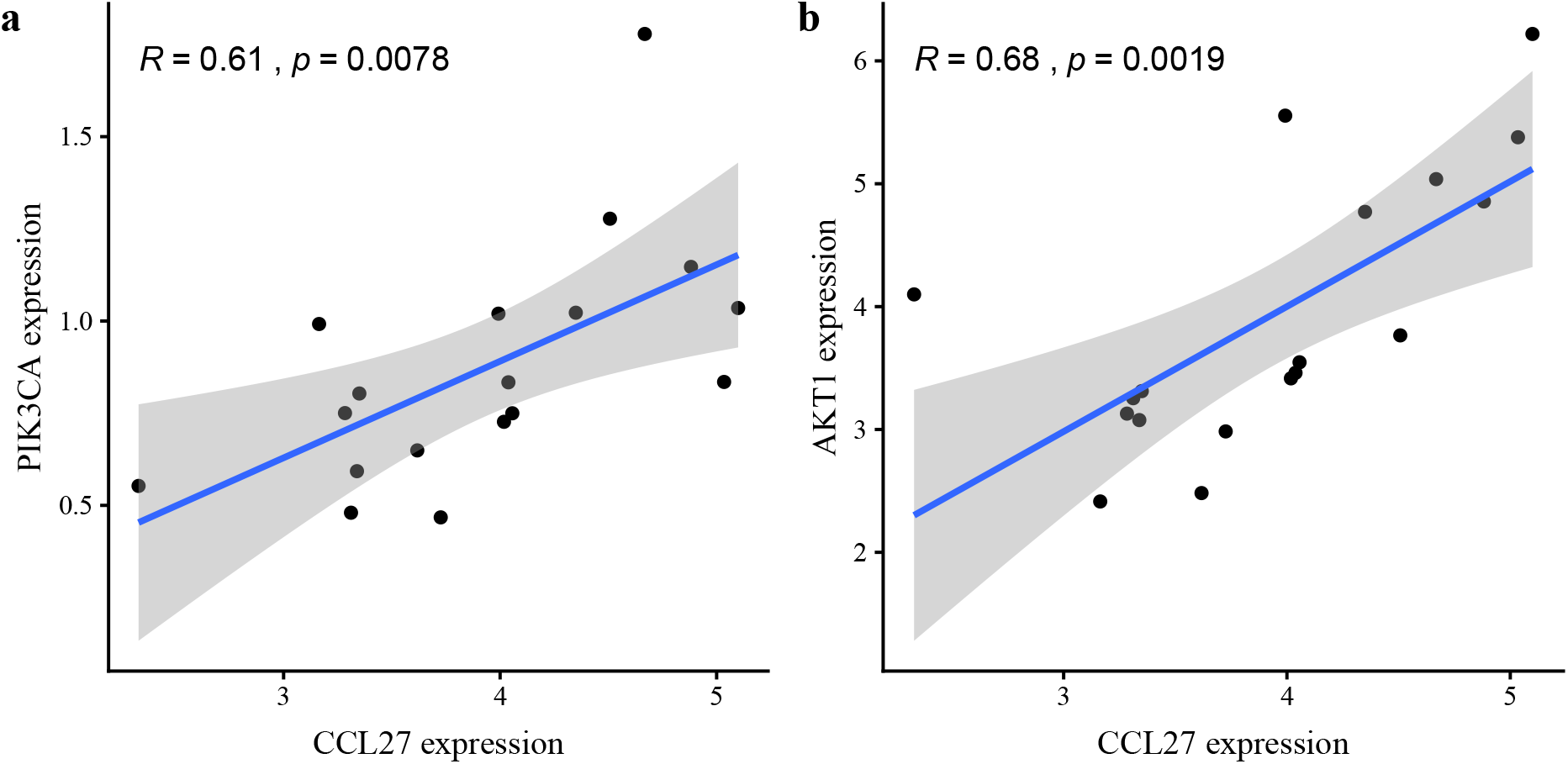
The *CCL27:CCR10* axis is upregulated in progressive samples and correlates with PIK/AKT expression. We compared ligand:receptor expression for each of 52 known cytokine:receptor pairs in 18 CIS lesions (n=10 progressive, 8 regressive). Only *CCL27:CCR10* was significantly different between progressive and regressive lesions (FDR 0.003; Figure 4). Progressive samples showed upregulated *CCL27* and downregulated *CCR10*. *CCL27* activation of *CCR10* has been shown to promote immune escape in mouse models, with the PIK/Akt pathway implicated as a potential mechanism. In CIS data, *CCL27* expression correlates with expression of both *PIK3CA* (a) and *AKT1* (b). Correlation coefficients are calculated using Pearson’s product moment.

**Extended Data Fig 9.**
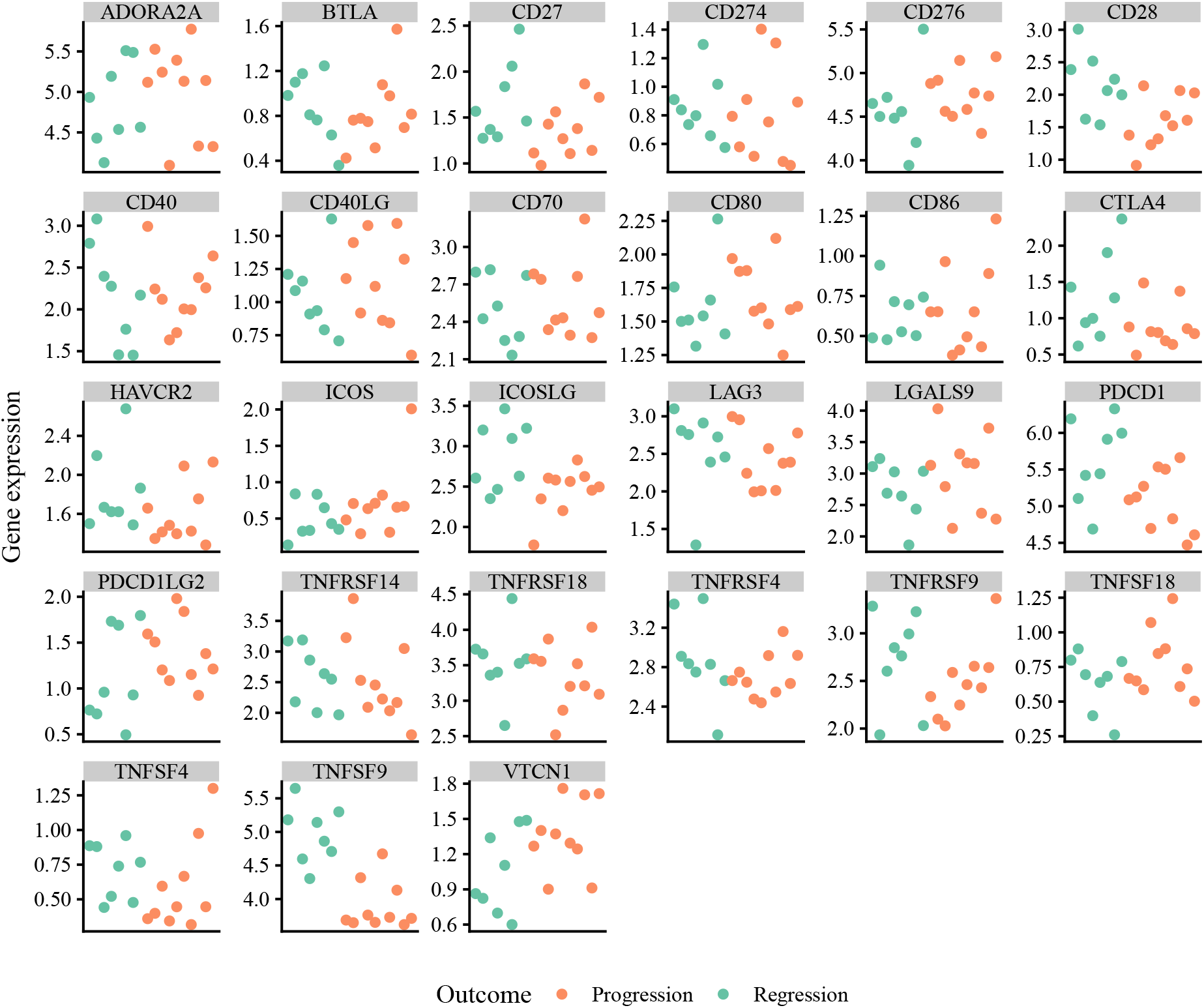
Comparisons of immune checkpoint molecules between progressive and regressive CIS samples. Here we show gene expression values of immune checkpoint molecules for each individual CIS lesion, showing both progressive (red; n=10) and regressive (blue; n=8). Although only *TNFSF9* reaches a significance threshold of FDR < 0.05 on differential expression analysis, other genes show outlier samples in the progressive group. Defects in these genes may be a critical immune escape mechanism in these outlier samples.

**Extended Data Fig 10.**
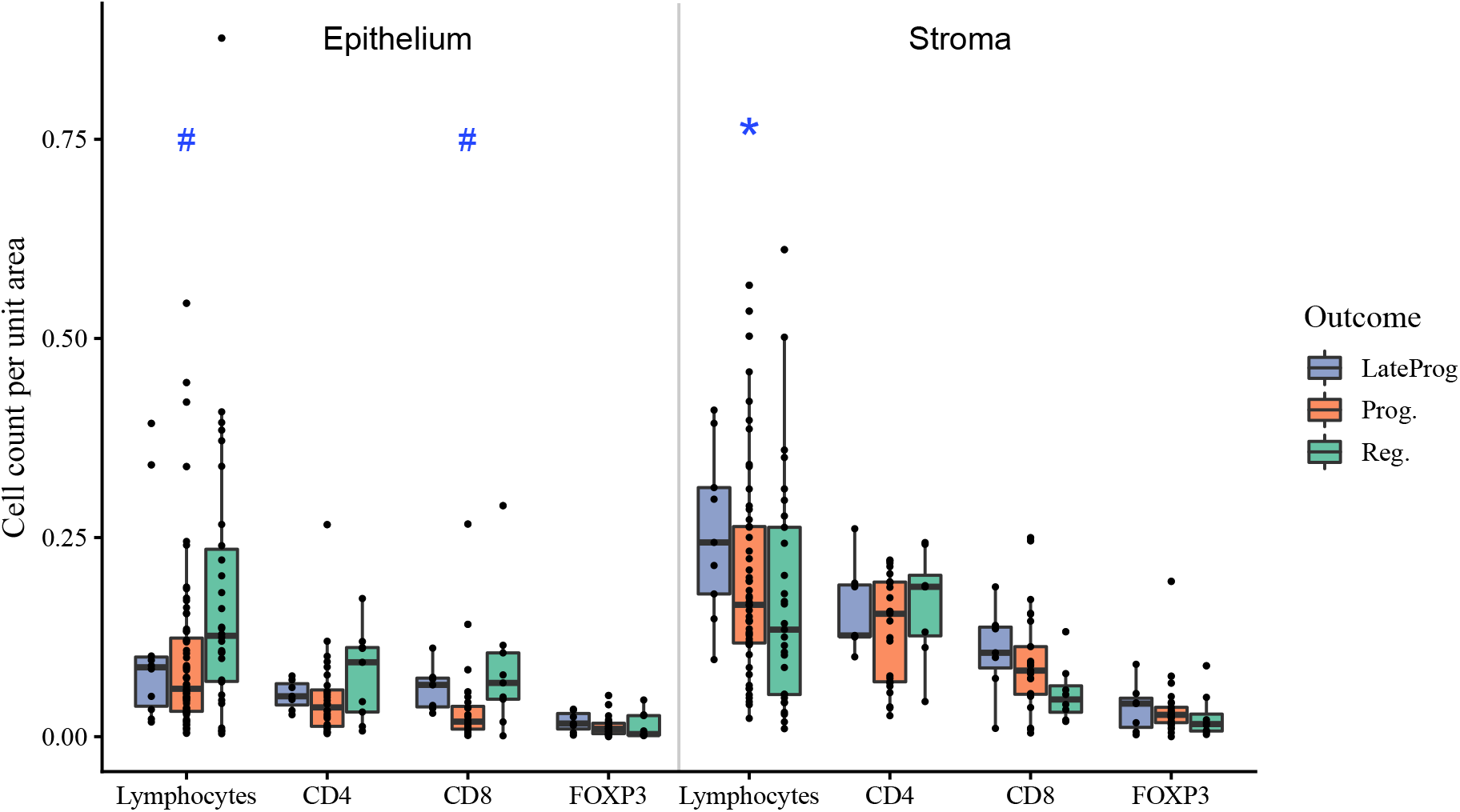
Of 53 lesions that met the clinical endpoint for regression – defined as a subsequent biopsy showing normal epithelium or low-grade dysplasia – 11 developed cancer later at the same site. These are termed ‘late progressive’ lesions. Combined quantitative immunohistochemistry data (n=44; 28 progressive, 16 regressive) with lymphocyte quantification from H&E images (n=116; 69 progressive, 47 regressive) are shown. We observe a similar trend of increased lymphocytes (p=0.06) and CD8+ cells (p=0.08) in regressive and late progressive samples compared to progressive. We also observe increased stromal lymphocytes in the late progressive group (p=0.02). Quoted p-values are calculated using ANOVA to reject the null hypothesis that all groups are equal, based on a linear mixed model to correct for multiple samples per patient; *<0.05, ^#^<0.1. Post-hoc pairwise comparisons using a Tukey HSD test were performed but sample size was insufficient to show significant results.

## Bibliography

1. A. G. Nicholson, L. J. Perry, P. M. Cury, P. Jackson, C. M. McCormick, B. Corrin, and A. U. Wells. Reproducibility of the WHO/IASLC grading system for pre-invasive squamous lesions of the bronchus: a study of inter-observer and intra-observer variation. Histopathology, 38 (3):202–8, 2001. ISSN 0309-0167 (Print) 0309-0167 (Linking).

2. Céline Mascaux, Mihaela Angelova, Angela Vasaturo, Jennifer Beane, Kahkeshan Hijazi, Geraldine Anthoine, Bénédicte Buttard, Françoise Rothe, Karen Willard-Gallo, Annick Haller, Vincent Ninane, Arsène Burny, Jean-Paul Sculier, Avi Spira, and Jérôme Galon. Immune evasion before tumour invasion in early lung squamous carcinogenesis. Nature, June 2019. ISSN 1476-4687. doi: 10.1038/s41586-019-1330-0.

3. Philip Jeremy George, Anindo K. Banerjee, Catherine A. Read, Caoihme O’Sullivan, Mary Falzon, Francesco Pezzella, Andrew G. Nicholson, Penny Shaw, Geoff Laurent, and Pamela H. Rabbitts. Surveillance for the detection of early lung cancer in patients with bronchial dysplasia. Thorax, 62(1):43–50, January 2007. ISSN 0040-6376, 1468-3296. doi: 10.1136/thx.2005.052191.

4. Martin Oft. IL-10: Master Switch from Tumor-Promoting Inflammation to Antitumor Immunity. Cancer Immunology Research, 2(3):194–199, March 2014. ISSN 2326-6066, 2326-6074. doi: 10.1158/2326-6066.CIR-13-0214.

5. Rachel Rosenthal, Elizabeth Larose Cadieux, Roberto Salgado, Maise Al Bakir, David A. Moore, Crispin T. Hiley, Tom Lund, Miljana Tanić, James L. Reading, Kroopa Joshi, Jake Y. Henry, Ehsan Ghorani, Gareth A. Wilson, Nicolai J. Birkbak, Mariam Jamal-Hanjani, Selvaraju Veeriah, Zoltan Szallasi, Sherene Loi, Matthew D. Hellmann, Andrew Feber, Benny Chain, Javier Herrero, Sergio A. Quezada, Jonas Demeulemeester, Peter Van Loo, Stephan Beck, Nicholas McGranahan, and Charles Swanton. Neoantigen-directed immune escape in lung cancer evolution. Nature, page 1, March 2019. ISSN 1476-4687. doi: 10.1038/s41586-019-1032-7.

6. Jennifer E. Beane, Sarah A. Mazzilli, Joshua D. Campbell, Grant Duclos, Kostyantyn Krysan, Christopher Moy, Catalina Perdomo, Michael Schaffer, Gang Liu, Sherry Zhang, Hanqiao Liu, Jessica Vick, Samjot S. Dhillon, Suso J. Platero, Steven M. Dubinett, Christopher Stevenson, Mary E. Reid, Marc E. Lenburg, and Avrum E. Spira. Molecular subtyping reveals immune alterations associated with progression of bronchial premalignant lesions. Nature Communications, 10(1):1856, April 2019. ISSN 2041-1723. doi: 10.1038/s41467-019-09834-2.

7. N. McGranahan, A. J. Furness, R. Rosenthal, S. Ramskov, R. Lyngaa, S. K. Saini, M. Jamal-Hanjani, G. A. Wilson, N. J. Birkbak, C. T. Hiley, T. B. Watkins, S. Shafi, N. Murugaesu, R. Mitter, A. U. Akarca, J. Linares, T. Marafioti, J. Y. Henry, E. M. Van Allen, D. Miao, B. Schilling, D. Schadendorf, L. A. Garraway, V. Makarov, N. A. Rizvi, A. Snyder, M. D. Hellmann, T. Merghoub, J. D. Wolchok, S. A. Shukla, C. J. Wu, K. S. Peggs, T. A. Chan, S. R. Hadrup, S. A. Quezada, and C. Swanton. Clonal neoantigens elicit T cell immunore-activity and sensitivity to immune checkpoint blockade. Science, 351(6280):1463–9, 2016. ISSN 1095-9203 (Electronic) 0036-8075 (Linking). doi: 10.1126/science.aaf1490.

8. V. H. Teixeira, C. P. Pipinikas, Adam Pennycuick, H. Lee-Six, Deepak Chandrasekharan, Jennifer Beane, T. J. Morris, Anna Karpathakis, A. Feber, C. E. Breeze, P Ntolios, R. E. Hynds, Mary Falzon, A. Capitanio, B. Carroll, P Durrenberger, G Hardavella, J. M. Brown, Andy G. Lynch, James Henry Royston Farmery, D. S. Paul, R Chambers, N. McGranahan, N. Navani, R. Thakrar, C. Swanton, S. Beck, J. P. George, C. Thirlwell, and S. Janes. Deciphering the genomic, epigenomic and transcriptomic landscapes of preinvasive lung cancer lesions. Nature Medicine, 2019. doi: 10.1038/s41591-018-0323-0.

9. Eszter Lakatos, Marc J. Williams, Ryan O. Schenck, William C. H. Cross, Jacob Househam, Benjamin Werner, Chandler Gatenbee, Mark Robertson-Tessi, Chris P. Barnes, Alexander R. A. Anderson, Andrea Sottoriva, and Trevor A. Graham. Evolutionary dynamics of neoantigens in growing tumours. bioRxiv, page 536433, January 2019. doi: 10.1101/536433.

10. N. McGranahan, R. Rosenthal, C. T. Hiley, A. J. Rowan, T. B. K. Watkins, G. A. Wilson, N. J. Birkbak, S. Veeriah, P. Van Loo, J. Herrero, C. Swanton, and T. RACERx Consortium. Allele-Specific HLA Loss and Immune Escape in Lung Cancer Evolution. Cell, 171(6):1259–1271 e11, 2017. ISSN 1097-4172 (Electronic) 0092-8674 (Linking). doi: 10.1016/j.cell.2017.10.001.

11. Balázs Győrffy, Giulia Bottai, Thomas Fleischer, Gyöngyi Munkácsy, Jan Budczies, Laura Paladini, Anne-Lise Børresen-Dale, Vessela N. Kristensen, and Libero Santarpia. Aberrant DNA methylation impacts gene expression and prognosis in breast cancer subtypes. International Journal of Cancer, 138(1):87–97, January 2016. ISSN 1097-0215. doi: 10.1002/ijc.29684.

12. Q. Ye, Y. Shen, X. Wang, J. Yang, F. Miao, C. Shen, and J. Zhang. Hypermethylation of HLA class I gene is associated with HLA class I down-regulation in human gastric cancer. Tissue Antigens, 75(1):30–39, January 2010. ISSN 1399-0039. doi: 10.1111/j.1399-0039.2009.01390.x.

13. The Cancer Genome Atlas Research Network. Comprehensive genomic characterization of squamous cell lung cancers. Nature, 489(7417):519–525, September 2012. ISSN 1476-4687. doi: 10.1038/nature11404.

14. Lei Wang, Zohreh Amoozgar, Jing Huang, Mohammad H. Saleh, Deyin Xing, Sandra Orsulic, and Michael S. Goldberg. Goldberg. Decitabine Enhances Lymphocyte Migration and Function and Synergizes with CTLA-4 Blockade in a Murine Ovarian Cancer Model. Cancer Immunology Research, 3(9):1030–1041, September 2015. ISSN 2326-6074. doi: 10.1158/2326-6066.CIR-15-0073.

15. Li-Xin Wang, Zhen-Yang Mei, Ji-Hao Zhou, Yu-Shi Yao, Yong-Hui Li, Yi-Han Xu, Jing-Xin Li, Xiao-Ning Gao, Min-Hang Zhou, Meng-Meng Jiang, Li Gao, Yi Ding, Xue-Chun Lu, Jin-Long Shi, Xu-Feng Luo, Jia Wang, Li-Li Wang, Chunfeng Qu, Xue-Feng Bai, and Li Yu. Low dose decitabine treatment induces CD80 expression in cancer cells and stimulates tumor specific cytotoxic T lymphocyte responses. PloS One, 8(5):e62924, 2013. ISSN 1932-6203. doi: 10.1371/journal.pone.0062924.

16. H. Yang, C. Bueso-Ramos, C. DiNardo, M. R. Estecio, M. Davanlou, Q.-R. Geng, Z. Fang, M. Nguyen, S. Pierce, Y. Wei, S. Parmar, J. Cortes, H. Kantarjian, and G. Garcia-Manero. Expression of PD-L1, PD-L2, PD-1 and CTLA4 in myelodysplastic syndromes is enhanced by treatment with hypomethylating agents. Leukemia, 28(6):1280–1288, June 2014. ISSN 1476-5551. doi: 10.1038/leu.2013.355.

17. Phase II Anti-PD1 Epigenetic Therapy Study in NSCLC. - Full Text View - ClinicalTrials.gov,.

18. Pembrolizumab (Immunotherapy Drug) in Combination With Guadecitabine and Mocetinostat (Epigenetic Drugs) for Patients With Advanced Lung Cancer. - Full Text View - ClinicalTrials.gov,.

19. Amel Saadi, Nicholas B. Shannon, Pierre Lao-Sirieix, Maria O’Donovan, Elaine Walker, Nicholas J. Clemons, James S. Hardwick, Chunsheng Zhang, Madhumita Das, Vicki Save, Marco Novelli, Frances Balkwill, and Rebecca C. Fitzgerald. Stromal genes discriminate preinvasive from invasive disease, predict >outcome, and highlight inflammatory pathways in digestive cancers. Proceedings of the National Academy of Sciences, 107(5):2177–2182, February 2010. ISSN 0027-8424, 1091-6490. doi: 10.1073/pnas.0909797107.

20. Rajani Ravi, Kimberly A. Noonan, Vui Pham, Rishi Bedi, Alex Zhavoronkov, Ivan V. Ozerov, Eugene Makarev, Artem V. Artemov, Piotr T. Wysocki, Ranee Mehra, Sridhar Nimmagadda, Luigi Marchionni, David Sidransky, Ivan M. Borrello, Evgeny Izumchenko, and Atul Bedi. Bifunctional immune checkpoint-targeted antibody-ligand traps that simultaneously disable TGFβ enhance the efficacy of cancer immunotherapy. Nature Communications, 9(1):741, February 2018. ISSN 2041-1723. doi: 10.1038/s41467-017-02696-6.

21. Mitsuo Yamauchi, Thomas H. Barker, Don L. Gibbons, and Jonathan M. Kurie. The fibrotic tumor stroma. The Journal of Clinical Investigation, 128(1):16–25, January 2018. ISSN 0021-9738. doi: 10.1172/JCI93554.

22. Sanjeev Mariathasan, Shannon J. Turley, Dorothee Nickles, Alessandra Castiglioni, Kobe Yuen, Yulei Wang, Edward E. Kadel Iii, Hartmut Koeppen, Jillian L. Astarita, Rafael Cubas, Suchit Jhunjhunwala, Romain Banchereau, Yagai Yang, Yinghui Guan, Cecile Chalouni, James Ziai, Yasin Şenbabaoğlu, Stephen Santoro, Daniel Sheinson, Jeffrey Hung, Jennifer M. Giltnane, Andrew A. Pierce, Kathryn Mesh, Steve Lianoglou, Johannes Riegler, Richard A. D. Carano, Pontus Eriksson, Mattias Höglund, Loan Somarriba, Daniel L. Halligan, Michiel S. van der Heijden, Yohann Loriot, Jonathan E. Rosenberg, Lawrence Fong, Ira Mellman, Daniel S. Chen, Marjorie Green, Christina Derleth, Gregg D. Fine, Priti S. Hegde, Richard Bourgon, and Thomas Powles. TGFβ attenuates tumour response to PD-L1 blockade by contributing to exclusion of T cells. Nature, 554(7693):544–548, February 2018. ISSN 1476-4687. doi: 10.1038/nature25501.

23. Min Zhao, Lei Kong, Yining Liu, and Hong Qu. dbEMT: an epithelial-mesenchymal transition associated gene resource. Scientific Reports, 5, June 2015. ISSN 2045-2322. doi: 10.1038/srep11459.

24. Yanyan Lou, Lixia Diao, Edwin Roger Parra Cuentas, Warren L. Denning, Limo Chen, You Hong Fan, Lauren A. Byers, Jing Wang, Vassiliki A. Papadimitrakopoulou, Carmen Behrens, Jaime Canales Rodriguez, Patrick Hwu, Ignacio I. Wistuba, John V. Heymach, and Don L. Gibbons. Epithelial-Mesenchymal Transition Is Associated with a Distinct Tumor Microenvironment Including Elevation of Inflammatory Signals and Multiple Immune Checkpoints in Lung Adenocarcinoma. Clinical Cancer Research: An Official Journal of the American Association for Cancer Research, 22(14):3630–3642, 2016. ISSN 1078-0432. doi: 10.1158/1078-0432.CCR-15-1434.

25. Albert Zlotnik, Osamu Yoshie, and Hisayuki Nomiyama. The chemokine and chemokine receptor superfamilies and their molecular evolution. Genome Biology, 7(12):243, 2006. ISSN 1465-6906. doi: 10.1186/gb-2006-7-12-243.

26. Takashi Murakami, Adela R. Cardones, Steven E. Finkelstein, Nicholas P. Restifo, Brenda A. Klaunberg, Frank O. Nestle, S. Sianna Castillo, Phillip A. Dennis, and Sam T. Hwang. Immune Evasion by Murine Melanoma Mediated through CC Chemokine Receptor-10. The Journal of Experimental Medicine, 198(9):1337–1347, November 2003. ISSN 0022-1007. doi: 10.1084/jem.20030593.

27. GTEx Consortium. Human genomics. The Genotype-Tissue Expression (GTEx) pilot analysis: multitissue gene regulation in humans. Science (New York, N.Y.), 348(6235):648–660, May 2015. ISSN 1095-9203. doi: 10.1126/science.1262110.

28. Luis Paz-Ares, Alexander Luft, David Vicente, Ali Tafreshi, Mahmut Gümüş, Julien Mazières, Barbara Hermes, Filiz Çay Şenler, Tibor Csőszi, Andrea Fülöp, Jerónimo Rodríguez-Cid, Jonathan Wilson, Shunichi Sugawara, Terufumi Kato, Ki Hyeong Lee, Ying Cheng, Silvia Novello, Balazs Halmos, Xiaodong Li, Gregory M. Lubiniecki, Bilal Piperdi, and Dariusz M. Kowalski. Pembrolizumab plus Chemotherapy for Squamous Non–Small-Cell Lung Cancer. New England Journal of Medicine, 379(21):2040–2051, November 2018. ISSN 0028-4793. doi: 10.1056/NEJMoa1810865.

29. Xinyue Qi, Fanlin Li, Yi Wu, Chen Cheng, Ping Han, Jieyi Wang, and Xuanming Yang. Optimization of 4-1bb antibody for cancer immunotherapy by balancing agonistic strength with FcγR affinity. Nature Communications, 10(1):2141, May 2019. ISSN 2041-1723. doi: 10.1038/s41467-019-10088-1.

30. I. Melero, W. W. Shuford, S. A. Newby, A. Aruffo, J. A. Ledbetter, K. E. Hellström, R. S. Mittler, and L. Chen. Monoclonal antibodies against the 4-1bb T-cell activation molecule eradicate established tumors. Nature Medicine, 3(6):682–685, June 1997. ISSN 1078-8956.

31. T. Bartkowiak and M. A. Curran. 4-1bb Agonists: Multi-Potent Potentiators of Tumor Immunity. Front Oncol, 5:117, 2015. ISSN 2234-943X (Print) 2234-943X (Linking). doi: 10.3389/fonc.2015.00117.

32. Neil H. Segal, Aiwu R. He, Toshihiko Doi, Ronald Levy, Shailender Bhatia, Michael J. Pishvaian, Rossano Cesari, Ying Chen, Craig B. Davis, Bo Huang, Aron D. Thall, and Ajay K. Gopal. Phase I Study of Single-Agent Utomilumab (PF-05082566), a 4-1bb/CD137 Ago-nist, in Patients with Advanced Cancer. Clinical Cancer Research: An Official Journal of the American Association for Cancer Research, 24(8):1816–1823, April 2018. ISSN 1078-0432. doi: 10.1158/1078-0432.CCR-17-1922.

33. C. P. Pipinikas, T. S. Kiropoulos, V. H. Teixeira, J. M. Brown, A. Varanou, M. Falzon, A. Capitanio, S. E. Bottoms, B. Carroll, N. Navani, F. McCaughan, J. P. George, A. Giangreco, N. A. Wright, S. A. McDonald, T. A. Graham, and S. M. Janes. Cell migration leads to spatially distinct but clonally related airway cancer precursors. Thorax, 69(6):548–57, 2014. ISSN 1468-3296 (Electronic) 0040-6376 (Linking). doi: 10.1136/thoraxjnl-2013-204198.

## Bibliography

1. Philip Jeremy George, Anindo K. Banerjee, Catherine A. Read, Caoihme O’Sullivan, Mary Falzon, Francesco Pezzella, Andrew G. Nicholson, Penny Shaw, Geoff Laurent, and Pamela H. Rabbitts. Surveillance for the detection of early lung cancer in patients with bronchial dysplasia. Thorax, 62(1):43–50, January 2007. ISSN 0040-6376, 1468-3296. doi: 10.1136/thx.2005.052191.

2. V. H. Teixeira, C. P. Pipinikas, Adam Pennycuick, H. Lee-Six, Deepak Chandrasekharan, Jennifer Beane, T. J. Morris, Anna Karpathakis, A. Feber, C. E. Breeze, P Ntolios, R. E. Hynds, Mary Falzon, A. Capitanio, B. Carroll, P Durrenberger, G Hardavella, J. M. Brown, Andy G. Lynch, James Henry Royston Farmery, D. S. Paul, R Chambers, N. McGranahan, N. Navani, R. Thakrar, C. Swanton, S. Beck, J. P. George, C. Thirlwell, and S. M. Janes. Deciphering the genomic, epigenomic and transcriptomic landscapes of preinvasive lung cancer lesions. Nature Medicine, 2019. doi: 10.1038/s41591-018-0323-0.

3. Laurent Gautier, Leslie Cope, Benjamin M. Bolstad, and Rafael A. Irizarry. affy—analysis of Affymetrix GeneChip data at the probe level. Bioinformatics, 20(3):307–315, February 2004. ISSN 1367-4803. doi: 10.1093/bioinformatics/btg405.

4. Tiffany J. Morris, Lee M. Butcher, Andrew Feber, Andrew E. Teschendorff, Ankur R. Chakravarthy, Tomasz K. Wojdacz, and Stephan Beck. ChAMP: 450k Chip Analysis Methylation Pipeline. Bioinformatics, 30(3):428–430, February 2014. ISSN 1367-4803. doi: 10.1093/bioinformatics/btt684.

5. D. Jones, K. M. Raine, H. Davies, P. S. Tarpey, A. P. Butler, J. W. Teague, S. Nik-Zainal, and P. J. Campbell. cgpCaVEManWrapper: Simple Execution of CaVEMan in Order to Detect Somatic Single Nucleotide Variants in NGS Data. Curr Protoc Bioinformatics, 56:15 10 1–15 10 18, 2016. ISSN 1934-340X (Electronic) 1934-3396 (Linking). doi: 10.1002/cpbi.20.

6. K. Ye, M. H. Schulz, Q. Long, R. Apweiler, and Z. Ning. Pindel: a pattern growth approach to detect break points of large deletions and medium sized insertions from paired-end short reads. Bioinformatics, 25(21):2865–71, 2009. ISSN 1367-4811 (Electronic) 1367-4803 (Linking). doi: 10.1093/bioinformatics/btp394.

7. K. M. Raine, J. Hinton, A. P. Butler, J. W. Teague, H. Davies, P. Tarpey, S. Nik-Zainal, and P. J. Campbell. cgpPindel: Identifying Somatically Acquired Insertion and Deletion Events from Paired End Sequencing. Curr Protoc Bioinformatics, 52:15 7 1–12, 2015. ISSN 1934-340X (Electronic) 1934-3396 (Linking). doi: 10.1002/0471250953.bi1507s52.

8. K. M. Raine, P. Van Loo, D. C. Wedge, D. Jones, A. Menzies, A. P. Butler, J. W. Teague, P. Tarpey, S. Nik-Zainal, and P. J. Campbell. ascatNgs: Identifying Somatically Acquired Copy-Number Alterations from Whole-Genome Sequencing Data. Curr Protoc Bioinformatics, 56:15 9 1–15 9 17, 2016. ISSN 1934-340X (Electronic) 1934-3396 (Linking). doi: 10.1002/cpbi.17.

9. Elli Papaemmanuil, Inmaculada Rapado, Yilong Li, Nicola E. Potter, David C. Wedge, Jose Tubio, Ludmil B. Alexandrov, Peter Van Loo, Susanna L. Cooke, John Marshall, Inigo Martincorena, Jonathan Hinton, Gunes Gundem, Frederik W. van Delft, Serena Nik-Zainal, David R. Jones, Manasa Ramakrishna, Ian Titley, Lucy Stebbings, Catherine Leroy, Andrew Menzies, John Gamble, Ben Robinson, Laura Mudie, Keiran Raine, Sarah O’Meara, Jon W. Teague, Adam P. Butler, Giovanni Cazzaniga, Andrea Biondi, Jan Zuna, Helena Kempski, Markus Muschen, Anthony M. Ford, Michael R. Stratton, Mel Greaves, and Peter J. Campbell. RAG-mediated recombination is the predominant driver of oncogenic rearrangement in *ETV6-RUNX1* acute lymphoblastic leukemia. Nature Genetics, 46(2):116–125, February 2014. ISSN 1546-1718. doi: 10.1038/ng.2874.

10. M. Kanehisa and S. Goto. KEGG: kyoto encyclopedia of genes and genomes. Nucleic Acids Res, 28(1):27–30, 2000. ISSN 0305-1048 (Print) 0305-1048 (Linking).

11. Wolfgang Huber, Vincent J. Carey, Robert Gentleman, Simon Anders, Marc Carlson, Benilton S. Carvalho, Hector Corrada Bravo, Sean Davis, Laurent Gatto, Thomas Girke, Raphael Gottardo, Florian Hahne, Kasper D. Hansen, Rafael A. Irizarry, Michael Lawrence, Michael I. Love, James MacDonald, Valerie Obenchain, Andrzej K. Oleś, Hervé Pagès, Alejandro Reyes, Paul Shannon, Gordon K. Smyth, Dan Tenenbaum, Levi Waldron, and Martin Morgan. Orchestrating high-throughput genomic analysis with Bioconductor. Nature Methods, 12(2):115–121, February 2015. ISSN 1548-7105. doi: o10.1038/nmeth.3252.

12. Douglas Bates, Martin Mächler, Ben Bolker, and Steve Walker. Fitting Linear Mixed-Effects Models Using lme4. Journal of Statistical Software, 67(1):1–48, October 2015. ISSN 1548-7660. doi: 10.18637/jss.v067.i01.

13. Applied Regression 3e.

14. M. E. Ritchie, B. Phipson, D. Wu, Y. Hu, C. W. Law, W. Shi, and G. K. Smyth. limma powers differential expression analyses for RNA-sequencing and microarray studies. Nucleic Acids Res, 43(7):e47, 2015. ISSN 1362-4962 (Electronic) 0305-1048 (Linking). doi: 10.1093/nar/gkv007.

15. Yoav Benjamini and Daniel Yekutieli. The control of the false discovery rate in multiple testing under dependency. The Annals of Statistics, 29(4):1165–1188, August 2001. ISSN 0090-5364, 2168-8966. doi: 10.1214/aos/1013699998.

16. Raivo Kolde. Pheatmap: pretty heatmaps. R package version, 61, 2012.

17. M. Jamal-Hanjani, G. A. Wilson, N. McGranahan, N. J. Birkbak, T. B. K. Watkins, S. Veeriah, S. Shafi, D. H. Johnson, R. Mitter, R. Rosenthal, M. Salm, S. Horswell, M. Escudero, N. Matthews, A. Rowan, T. Chambers, D. A. Moore, S. Turajlic, H. Xu, S. M. Lee, M. D. Forster, T. Ahmad, C. T. Hiley, C. Abbosh, M. Falzon, E. Borg, T. Marafioti, D. Lawrence, M. Hayward, S. Kolvekar, N. Panagiotopoulos, S. M. Janes, R. Thakrar, A. Ahmed, F. Blackhall, Y. Summers, R. Shah, L. Joseph, A. M. Quinn, P. A. Crosbie, B. Naidu, G. Middleton, G. Langman, S. Trotter, M. Nicolson, H. Remmen, K. Kerr, M. Chetty, L. Gomersall, D. A. Fennell, A. Nakas, S. Rathinam, G. Anand, S. Khan, P. Russell, V. Ezhil, B. Ismail, M. Irvin-Sellers, V. Prakash, J. F. Lester, M. Kornaszewska, R. Attanoos, H. Adams, H. Davies, S. Dentro, P. Taniere, B. O’Sullivan, H. L. Lowe, J. A. Hartley, N. Iles, H. Bell, Y. Ngai, J. A. Shaw, J. Herrero, Z. Szallasi, R. F. Schwarz, A. Stewart, S. A. Quezada, J. Le Quesne, P. Van Loo, C. Dive, A. Hackshaw, C. Swanton, and T. RACERx Consortium. Tracking the Evolution of Non-Small-Cell Lung Cancer. N Engl J Med, 376(22):2109–2121, 2017. ISSN 1533-4406 (Electronic) 0028-4793 (Linking). doi: 10.1056/NEJMoa1616288.

18. K. Sirinukunwattana, S. E. A. Raza, Y. Tsang, D. R. J. Snead, I. A. Cree, and N. M. Rajpoot. Locality Sensitive Deep Learning for Detection and Classification of Nuclei in Routine Colon Cancer Histology Images. IEEE Transactions on Medical Imaging, 35(5):1196–1206, May 2016. ISSN 0278-0062. doi: 10.1109/TMI.2016.2525803.

19. David Allan Moore, Marco Sereno, Madhumita Das, Juvenal Dario Baena Acevedo, Samantha Sinnadurai, Claire Smith, Abi McSweeney, Xiaoyu Su, Leah Officer, Carolyn Jones, Kate Dudek, David Guttery, Phillipe Taniere, Ruth V. Spriggs, and John Le Quesne. In situ growth in early lung adenocarcinoma may represent precursor growth or invasive clone outgrowth—a clinically relevant distinction. Modern Pathology, page 1, April 2019. ISSN 1530-0285. doi: 10.1038/s41379-019-0257-1.

20. Ayse U. Akarca, Vishvesh H. Shende, Alan D. Ramsay, Tim Diss, Maria Pane-Foix, Hasan Rizvi, Maria R. Calaminici, Thomas M. Grogan, David Linch, and Teresa Marafioti. BRAF V600e mutation-specific antibody, a sensitive diagnostic marker revealing minimal residual disease in hairy cell leukaemia. British Journal of Haematology, 162(6):848–851, September 2013. ISSN 1365-2141. doi: 10.1111/bjh.12429.

21. Teresa Marafioti, Margaret Jones, Fabio Facchetti, Tim C. Diss, Ming-Qing Du, Peter G. Isaacson, Michela Pozzobon, Stefano A. Pileri, Amanda J. Strickson, Soo-Yong Tan, Fiona Watkins, and David Y. Mason. Phenotype and genotype of interfollicular large B cells, a subpopulation of lymphocytes often with dendritic morphology. Blood, 102(8):2868–2876, October 2003. ISSN 0006-4971. doi: 10.1182/blood-2003-03-0692.

22. András Szolek, Benjamin Schubert, Christopher Mohr, Marc Sturm, Magdalena Feldhahn, and Oliver Kohlbacher. OptiType: precision HLA typing from next-generation sequencing data. Bioinformatics, 30(23):3310–3316, December 2014. ISSN 1367-4803. doi: 10.1093/bioinformatics/btu548.

23. Massimo Andreatta and Morten Nielsen. Gapped sequence alignment using artificial neural networks: application to the MHC class I system. Bioinformatics (Oxford, England), 32(4): 511–517, February 2016. ISSN 1367-4811. doi: 10.1093/bioinformatics/btv639.

24. Morten Nielsen, Claus Lundegaard, Peder Worning, Sanne Lise Lauemøller, Kasper Lamberth, Søren Buus, Søren Brunak, and Ole Lund. Reliable prediction of T-cell epitopes using neural networks with novel sequence representations. Protein Science: A Publication of the Protein Society, 12(5):1007–1017, May 2003. ISSN 0961-8368. doi: 10.1110/ps.0239403.

25. N. McGranahan, R. Rosenthal, C. T. Hiley, A. J. Rowan, T. B. K. Watkins, G. A. Wilson, N. J. Birkbak, S. Veeriah, P. Van Loo, J. Herrero, C. Swanton, and T. RACERx Consortium. Allele-Specific HLA Loss and Immune Escape in Lung Cancer Evolution. Cell, 171(6):1259–1271 e11, 2017. ISSN 1097-4172 (Electronic) 0092-8674 (Linking). doi: 10.1016/j.cell.2017.10.001.

26. P. Danaher, S. Warren, L. Dennis, L. D’Amico, A. White, M. L. Disis, M. A. Geller, K. Odunsi, J. Beechem, and S. P. Fling. Gene expression markers of Tumor Infiltrating Leukocytes. J Immunother Cancer, 5:18, 2017. ISSN 2051-1426 (Electronic) 2051-1426 (Linking). doi: 10.1186/s40425-017-0215-8.

27. Rachel Rosenthal, Elizabeth Larose Cadieux, Roberto Salgado, Maise Al Bakir, David A. Moore, Crispin T. Hiley, Tom Lund, Miljana Tanić, James L. Reading, Kroopa Joshi, Jake Y. Henry, Ehsan Ghorani, Gareth A. Wilson, Nicolai J. Birkbak, Mariam Jamal-Hanjani, Selvaraju Veeriah, Zoltan Szallasi, Sherene Loi, Matthew D. Hellmann, Andrew Feber, Benny Chain, Javier Herrero, Sergio A. Quezada, Jonas Demeulemeester, Peter Van Loo, Stephan Beck, Nicholas McGranahan, and Charles Swanton. Neoantigen-directed immune escape in lung cancer evolution. Nature, page 1, March 2019. ISSN 1476-4687. doi: 10.1038/s41586-019-1032-7.

28. Ankur Chakravarthy, Andrew Furness, Kroopa Joshi, Ehsan Ghorani, Kirsty Ford, Matthew J. Ward, Emma V. King, Matt Lechner, Teresa Marafioti, Sergio A. Quezada, Gareth J. Thomas, Andrew Feber, and Tim R. Fenton. Pan-cancer deconvolution of tumour composition using DNA methylation. Nature Communications, 9(1):3220, August 2018. ISSN 2041-1723. doi: 10.1038/s41467-018-05570-1.

29. Aaron M. Newman, Chih Long Liu, Michael R. Stratton, Andrew J. Gentles, Weiguo Feng, Yue Xu, Chuong D. Hoang, Maximilian Diehn, and Ash A. Alizadeh. Robust enumeration of cell subsets from tissue expression profiles. Nature Methods, 12(5):453–457, May 2015. ISSN 1548-7105. doi: 10.1038/nmeth.3337.

